# High temporal frequency light response in mouse retina requires FAT3 signaling in bipolar cells

**DOI:** 10.1101/2023.11.02.565326

**Authors:** Evelyn C. Avilés, Sean K. Wang, Sarina Patel, Shuxiang Shi, Lucas Lin, Vladimir J. Kefalov, Lisa V. Goodrich, Constance L. Cepko, Yunlu Xue

**Affiliations:** Department of Neurobiology, Blavatnik Institute, Harvard Medical School, Boston, MA 02115; Facultad de Ciencias Biologicas, Pontificia Universidad Catolica de Chile, Santiago, Chile; Departments of Genetics and Ophthalmology, Harvard Medical School, Boston, MA 02115; Howard Hughes Medical Institute, Boston, MA 02115; Lingang Laboratory, Shanghai, China, 200031; School of Life Science and Technology, ShanghaiTech University, Shanghai, China, 201210; Gavin Herbert Eye Institute & Center for Translational Vision Research, University of California, Irvine, CA 92697

**Keywords:** high frequency vision, retinal physiology, FAT cadherins, bipolar cells, GRIK1

## Abstract

Vision is initiated by the reception of light by photoreceptors and subsequent processing via downstream retinal neurons. Proper cellular organization depends on the multi-functional tissue polarity protein FAT3, which is required for amacrine cell connectivity and retinal lamination. Here we investigated the retinal function of *Fat3* mutant mice and found decreases in physiological and perceptual responses to high frequency flashes. These defects did not correlate with abnormal amacrine cell wiring, pointing instead to a role in bipolar cell subtypes that also express FAT3. The role of FAT3 in the response to high temporal frequency flashes depends upon its ability to transduce an intracellular signal. Mechanistically, FAT3 binds to the synaptic protein PTPσ, intracellularly, and is required to localize GRIK1 to OFF-cone bipolar cell synapses with cone photoreceptors. These findings expand the repertoire of FAT3’s functions and reveal its importance in bipolar cells for high frequency light response.

## Introduction

The remarkable ability to detect light over a wide range of intensities and frequencies is accomplished by highly specialized cell types within the retina. Retinal circuits encode signals originating in the photoreceptors, which synapse with bipolar cells (BCs). BCs are classified as ON or OFF depending upon their response to photoreceptor signals, thereby setting the stage for downstream processing events. Critical to these transformations are the >80 types of retinal interneurons^1–4^, which are organized into laminae with the cell bodies of the horizontal, BCs, and amacrine cells (ACs) located in the inner nuclear layer (INL). In the adjacent outer nuclear layer (ONL) reside the cell bodies of the photoreceptors. Retinal interneurons form synaptic connections in two layers: in the outer plexiform layer (OPL) among BCs, horizontal cells and photoreceptors, and in the inner plexiform layer (IPL) among BCs, ACs, and the output neurons of the retina, the retinal ganglion cells (RGCs) (**Figure 1A**). Despite the stereotyped and conserved organization of retinal neurons and their connections, it is unclear how important the lamination is for the sense of vision. Also unknown is how the organization of most retinal neurons mediate specific visual responses, e.g. which cell types and connections are needed to process and transmit high frequency signals^5^.

**Figure 1:**
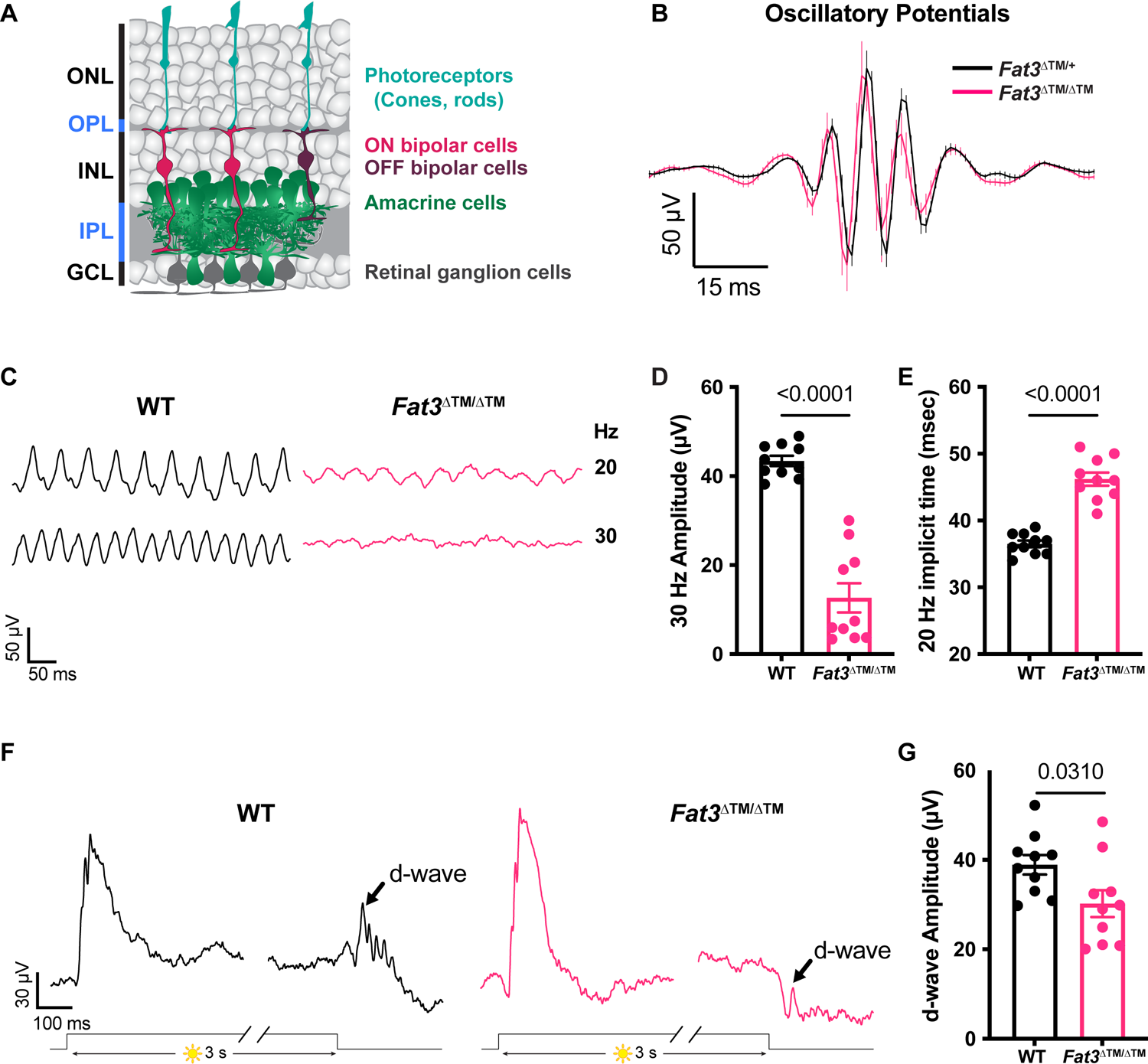
Flicker ERG and vision at high frequency and step ERG of *Fat3*-deficient mice. A. Schematic representation of retinal layers and their neurons. B. Averaged Oscillatory potentials of *Fat3*^ΔTM/+^ (n=8) and *Fat3*^ΔTM/ΔTM^ (n=10) eyes at scotopic condition elicited by 0.1 cd s/m^2^ flashes (same as Figure S1B). C. Representative flicker ERG raw traces of WT control and *Fat3*^ΔTM/ΔTM^ eyes elicited by 3.162 cd s/m^2^ flashes at 20 and 30 Hz frequencies. D. Flicker ERG amplitude at 30 Hz for WT control (n=10) and *Fat3*^ΔTM/ΔTM^ (n=10) eyes. Unpaired two-tailed Student’s t test. E. Flicker ERG implicit time (1^st^ peak) at 20 Hz for WT control (n=10) and *Fat3*^ΔTM/ΔTM^ (n=10) eyes. Unpaired two-tailed Student’s t test. F. Representative step ERG raw traces of WT (n=10) control and *Fat3*^ΔTM/ΔTM^ (n=10) eyes elicited by a 3-second step light at 1000 cd/m^2^ intensity. G. Statistics of step ERG amplitudes (b-wave, d-wave and b: d ratio) of WT (n=10) control and *Fat3*^ΔTM/ΔTM^ (n=10) eyes elicited by a 3-second step of light at 1000 cd/m^2^ intensity. Unpaired two-tailed Student’s t test.

Several features of retinal organization depend upon FAT3, a tissue polarity protein^6–8^. FAT cadherins are transmembrane receptors that can sense cell position in the environment *via* their huge extracellular domains. They induce appropriate changes in cell morphology *via* their intracellular domains, thereby creating cellular asymmetries that are aligned across the tissue. In *Fat3* mutant retinas, many ACs migrate to ectopic locations in the IPL and the ganglion cell layer (GCL). They also fail to retract their trailing neurites, which go on to form ectopic synapses in two misplaced plexiform layers, one in the INL (the outer misplaced plexiform layer, OMPL) and one below the GCL (the inner misplaced plexiform layer, IMPL)^6–8^. These ectopic layers contain synapses between ACs and between ACs and rod BCs, which do not express FAT3^2^, but seem to “follow” their AC partners to abnormal locations^7^. FAT3, which is localized to AC dendrites in the IPL, mediates these effects by localizing cytoskeletal effectors needed for migration and retraction^6^. The FAT3 intracellular domain also binds a variety of synaptic proteins^6^ and is therefore poised to control synapse localization or function independent of its effects on cell morphology. A direct functional role at the synapse has not been described, and it has not been clear how the loss of FAT3 impacts vision.

Here, we used retinal physiology and behavioral analyses to investigate the effects of FAT3 disruptions on retinal function. We discovered an additional function for FAT3 at the cone-BC synapse that impacts transmission of high temporal frequency signals. Basic light responses of retinal neurons were found to be preserved in *Fat3* mutant mice despite the highly abnormal pattern of lamination. However, overall retinal responses to 30 Hz flashes were severely reduced in amplitude and many *Fat3* mutants behaved as if they see constant illumination. Analysis of a series of conditional *Fat3* knock-out mice revealed that this phenotype is not due to altered AC position or connectivity. Instead, we found that the OFF-cone pathway is abnormal, as revealed by an *in vivo* electroretinogram (ERG) protocol that we developed. Further, we show that the FAT3 intracellular domain binds to the synaptic protein PTPσ, and that in *Fat3* mutants, both PTPσ and the glutamate receptor subunit GRIK1 are present at reduced levels in the ribbon synapses that link cones to OFF-cone BCs. Thus, FAT3 is required to set up and/or maintain a properly organized and functional synapse between photoreceptors and BCs and the transmission of high frequency signals, highlighting its multiple and versatile roles in the development and function of the retina.

## Results

### Global loss of *Fat3* affects the retinal response to high temporal frequency light

In *Fat3*^ΔTM/ΔTM^ mutant mice, which lack a membrane localized form of FAT3, retinal lamination is strongly disrupted, with changes in the position of ACs and their synapses with other ACs, as well as with other retinal cell types^7^. The abnormal lamination can also be seen by optical coherence tomography (OCT) imaging in animals *in vivo* (**Figure S1A**). To assess the functional consequences of this change in retinal cell organization, we used the electroretinogram (ERG), a common way to measure electrical changes in the retina in response to light. By altering the stimulus, it is possible to reveal the contributions of specific retinal pathways. For instance, signaling through the rod pathway is assayed by performing the ERG after dark-adaptation and under scotopic conditions, using dim flashes that elicit little response from the cone pathway. Conversely, photopic ERG tests isolate the cone pathway by light-adapting the eyes and using a background light that saturates rod phototransduction. In an ERG waveform, the a-wave originates from photoreceptors and is followed by the b-wave, which reflects the activity of rod bipolar cells (RBC) and/or ON-types of cone BCs (ON-CBCs). By flashing the stimulus on and off with increasing frequency, it is possible to determine how well photoreceptors and BCs are able to resolve temporal differences in the visual stimulus. As the frequency of the flash increases, OFF-CBCs are hypothesized to dominate the response^9^. Additionally, the ERG d-wave, which emerges when turning off a light step that lasts for a few seconds^10^, is thought to represent mainly OFF-CBC activity^11^.

Conventional ERG assays under scotopic and photopic conditions revealed no obvious difference in *Fat3*^ΔTM/ΔTM^ vs. *Fat3*^ΔTM/+^ littermates, indicating that overall signaling through rod and cone pathways was preserved (**Figure S1B-E**). Likewise, the oscillatory potentials between the a- and b-waves, which are thought to reflect the activity of ACs and/or RGCs^12^, showed no dramatic change in the number, amplitude or timing of peaks (**Figure 1B**). Thus, *Fat3*^ΔTM/ΔTM^ mutant mice can detect standard light stimuli despite abnormal retinal lamination.

Although previous work focused on its role in ACs^6,7^, *Fat3* is also expressed in RGCs and some subtypes of BCs, especially OFF-CBCs^2^. To determine whether *Fat3* is required for the proper response to high temporal frequency light, we recorded ERGs in response to lights that flicker in the high frequency range (i.e. over 15 Hz^9^). Flicker ERG responses in this range were hypothesized to be dominated by the OFF-pathway^9^. We observed a significant decrease in the amplitude of responses to lights flickering at 30 Hz in *Fat3*^ΔTM/ΔTM^ compared to their *Fat3*^ΔTM/+^ littermates (**Figure 1C,D**). Additionally, the timing of the first peak in response to 20 Hz stimulation (“the implicit time”) was delayed (**Figure 1E**). The 30 Hz implicit time was not measured, as many *Fat3* mutant mice presented no response at this frequency. These results suggested that the transmission of high temporal frequency light responses was impaired, presumably in BCs, where the signal to the flicker ERG likely originates^9^. To determine if this physiological deficit in the retina had perceptual consequences, behavioral assays were conducted to examine if the mutant mice can perceive high frequency flickering light, using contextual and vision-cued fear conditioning tests^13,14^. Normally, fear conditioned animals increase the time of “freezing” if they have been conditioned to associate an unpleasant stimulus with an environmental cue, in this case, 33 Hz light (**Figure S1F,G**, **Supplementary movies**). In contrast to *Fat3*^ΔTM/+^ mice (n=8 animals), some *Fat3*^ΔTM/ΔTM^ mice (5 out of 9 animals) failed to show a freezing response when switching from a static light to a 33 Hz flicker (**Figure S1G**), similar to the 30 Hz flicker ERG results (6 out of 10 *Fat3*^ΔTM/ΔTM^ eyes presented amplitudes smaller than the cut-off level of 20 μV, **Figure 1D**). The lack of response was not due to an inability to form fear memories, as all *Fat3*^ΔTM/ΔTM^ and *Fat3*^ΔTM/+^ animals froze less when switching from an unpleasant olfactory and tactile context (group “context”) to a novel and safe context within the group with static light (group “static light”) (**Figure S1G**). Mutant mice also performed like the wildtype (WT) controls in an optomotor behavioral assay, which measures spatial discrimination (**Figure S1H**).

Responses to different frequencies of light are used to study the temporal properties of vision at photoreceptor, brain, or psychophysical levels^15^. The flicker ERG at high temporal frequency has been proposed to depend on OFF-CBC function^9^. Usually, OFF-CBC function is probed using patch-clamping of single cells or assessed at the population level by examination of the d-wave of the ERG. However, to our knowledge, there has not been an ERG protocol to measure the d-wave from mice *in vivo*. To address this unmet need, we designed an *in vivo* ERG protocol for mice based on a study of the *ex vivo* ERG response of amphibian retinas. This assay measures the retinal voltage change, the d-wave, after turning off a long step of light (i.e. light-off)^10^, which is called a “step ERG”. Using this newly developed *in vivo* protocol on mice, we found that WT mice had a strong d-wave with robust oscillatory potentials at the end of a three-second exposure to a 1,000 cd/m^2^ step of light (**Figure 1F,G**). By contrast, in *Fat3*^ΔTM/ΔTM^ mice, the d-wave amplitude was decreased, and the d-wave related oscillatory potentials were absent (**Figure 1F,G**). Thus, responses to both high frequency flickering light and light-off stimuli showed that *Fat3*^ΔTM/ΔTM^ mice have deficits in processing specific types of visual stimuli that have been associated with OFF-CBCs.

To test the impact of an ectopic plexiform layer and other FAT3-dependent AC properties on retinal physiology, *Fat3* was deleted specifically from ACs using *Ptf1a*^CRE7,16^ and a *Fat3* floxed allele (**Figure 2A,B**). In these conditional knock-out (cKO) mutants, ACs migrate normally, but do not retract their trailing processes, leading to the formation of an OMPL^7^. We found that the 30 Hz flicker ERG amplitude and 20 Hz implicit time were normal in *Ptf1a*^CRE^ *Fat3*cKO mice compared to littermate *Ptf1a*^CRE^ control mice (**Figure 2C-E**), indicating that the high frequency light response defects were not secondary either to the presence of *Fat3* mutant ACs or the mild lamination defects they caused. This raised the possibility that the highly specific physiological deficits seen in *Fat3*^ΔTM/ΔTM^ mice are due instead to altered BC function.

**Figure 2:**
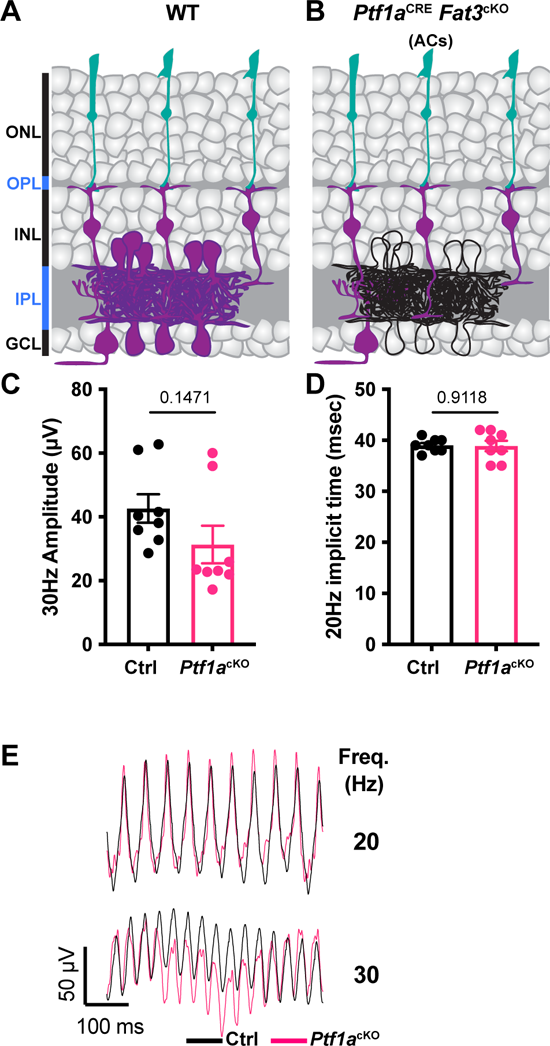
Flicker ERG at high frequency of *Ptf1a*^CRE^ conditional *Fat3* mice. A. Schematic representation of cell classes that express FAT3 in wild type tissue. Cells that express FAT3 are represented in magenta. B. Schematic representation of cell classes, i.e. ACs, that lose *Fat3* expression in a *Ptf1a*^CRE^ cKO, shown in black outlines. C. Flicker ERG amplitude at 30 Hz for the *Ptf1a*^CRE^ *Fat3*cKO condition. Control genotypes are *Ptf1a*^CRE^;*Fat3*^fl/+^ (n=8 eyes) and *Ptf1a*^cKO^ genotypes are *Ptf1a*^CRE^;*Fat3*^fl/ΔTM^ (n=8 eyes). Unpaired two-tailed Student’s t test. D. Flicker ERG implicit time at 20 Hz for *Ptf1a*^CRE^;*Fat3*^fl/+^ (control, n=8 eyes) and *Ptf1a*^CRE^;*Fat3*^fl/ΔTM^ (*Ptf1a*^cKO^, n=8 eyes). Unpaired two-tailed Student’s t test. E. Representative flicker ERG raw traces for *Ptf1a*^CRE^;*Fat3*^fl/+^ (control, n=8 eyes) and *Ptf1a*^CRE^;*Fat3*^fl/ΔTM^ (*Ptf1a*^cKO^, n=8 eyes).

Consistent with previous single cell RNA sequencing data showing its expression in CBC type 1A, 1B, 2 and 3A^2^, we confirmed that *Fat3* mRNA is present in BCs by *in situ* hybridization (**Figure 3A-C**). Additionally, FAT3 protein localizes not only to the IPL but also to the OPL, where BC dendrites form synapses with photoreceptors (**Figure 3D**). *In situ* hybridization confirmed co-expression of *Fat3* with the OFF-CBC marker *Grik1,* which encodes an ionotropic glutamate receptor^2,17^(**Figure 3F-I**). *Fat3* is also expressed in some *Grik1*-negative CBCs, which are positive for the ON-CBC marker *Grm6*, though to a less degree (**Figure 3I, S2**). This expression pattern is in line with FAT3 having a role on retinal functions associated with OFF-CBC pathways, as was suggested by the flicker and step ERG responses.

**Figure 3:**
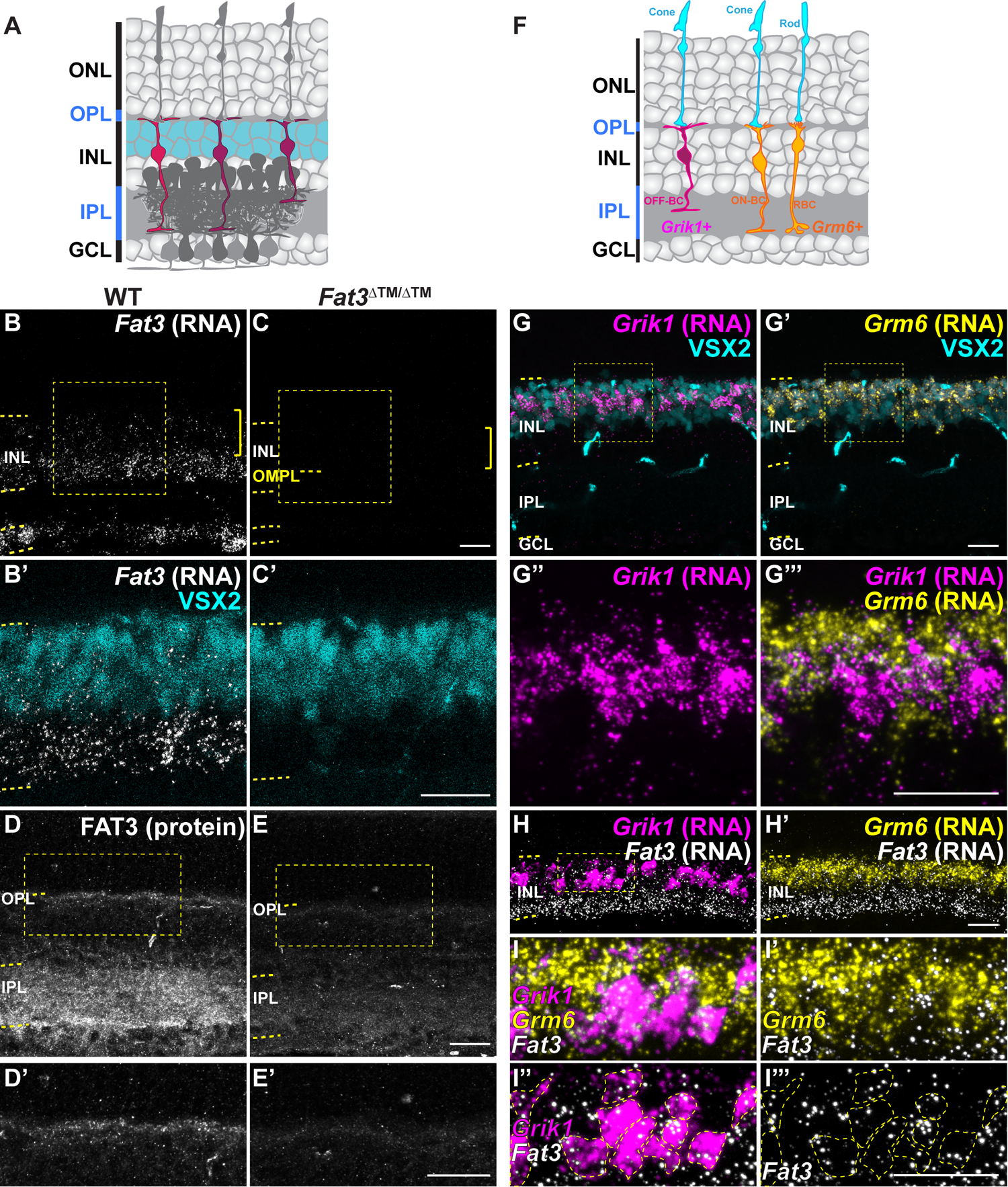
*Fat3* RNA is enriched in OFF-cone bipolar cells. A. Schematic representation of the retina highlighting the position of bipolar cell bodies, as stained by VSX2. B. *in situ* hybridization for *Fat3* RNA in WT P22 retinas. C. *in situ* hybridization for *Fat3* RNA in *Fat3*^ΔTM/ΔTM^ P22 retinal tissue. In B and C the RNA puncta are shown in white and the yellow brackets indicate the area of VSX2+ cell bodies. Yellow dashed lines demark the inner nuclear layer (INL) and the outer misplaced plexiform layer (OMPL) in *Fat3*^ΔTM/ΔTM^ tissue. The squares demark the insets seen in B’ and C’ at higher magnification. VSX2 protein is seen in cyan. D. Hybridization Chain Reaction-Immunohistochemistry (HCR-IHC) of FAT3 in wild type retinas. Inset demarked in a yellow box in d is shown at higher magnification in D’. E. HCR-IHC of FAT3 in *Fat3*^ΔTM/ΔTM^ mutant retinas. Inset demarked in a yellow box in e is shown at higher magnification in E’. F. Schematic representation of *Grik1* and *Grm6* RNA enrichment in bipolar cells, according to data in Figure S2. G. *in situ* hybridization of *Grik1* RNA (magenta) and Grm6 RNA (yellow, G’) with immunostaining for VSX2 (cyan). The insets in G and G’ are shown at a higher magnification in G’’ and G’’’. H. Triple *in situ* hybridization to *Fat3*, *Grik1* and *Grm6* RNA. Inset in H is seen at higher magnification in I-I’’’. I. Higher magnification of inset shown in H. Yellow dashed lines in I’’ and I’’’ demark *Grik1* RNA+ cell bodies. *Fat3* RNA (white) is shown together with *Grm6* RNA in I’, with *Grik1* RNA in I’’ and alone in I’’’. Scale bars: 20µm.

### Retinal lamination defects do not disrupt responses to flickering stimuli

We next asked how the presence of FAT3 in other retinal neurons influences retinal organization and contributes to the impaired high frequency light response seen in *Fat3*^ΔTM/ΔTM^ mutant mice. To survey other FAT3+ cells for possible effects on the high frequency light response, we used the *Isl1*^CRE^ mouse line to drive recombination in all ON-CBCs, starburst ACs, and RGCs, but not in OFF-CBCs^18^ (**Figure 4A-H**). Although *Isl1*^CRE^ *Fat3*cKO mice exhibited all the cellular phenotypes previously described in the retina of *Fat3*^ΔTM/ΔTM^ mice (**Figure 4B-H**), the 30 Hz flicker ERG amplitude and 20 Hz implicit time were normal compared to littermate *Isl1*^CRE^ control mice (**Figure 4I-K**).

**Figure 4:**
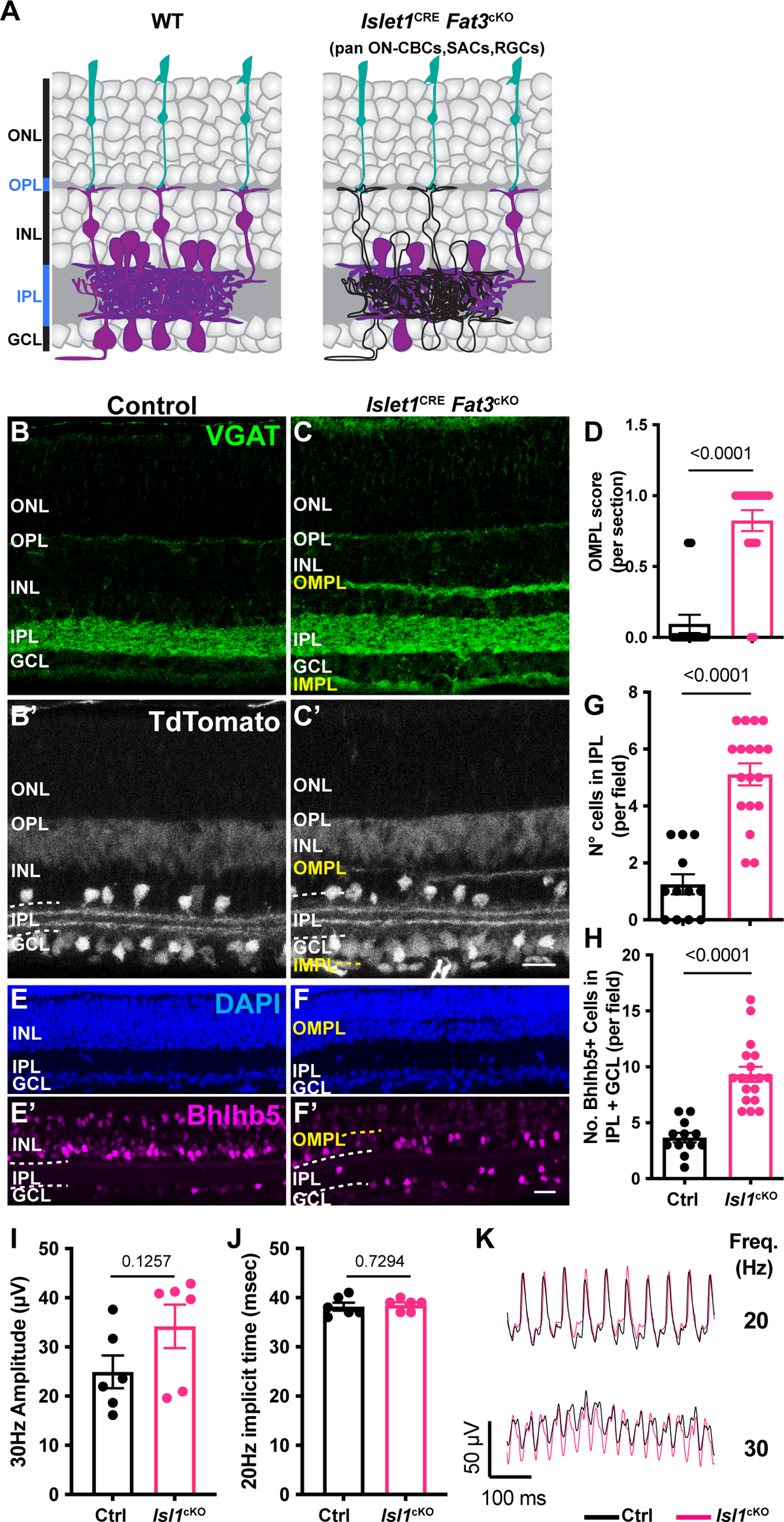
Flicker ERG at high frequency of *Ptf1a*^CRE^ and *Isl1*^CRE^ conditional *Fat3* mice. A. Schematic representation of cell classes, i.e. starburst ACs, RGCs and ON-CBCs, that lose *Fat3* expression in an *Isl1*^CRE^ cKO, shown in black outlines compared to magenta cells seen in WT controls. B. VGAT immunostaining for control *Isl1*^CRE/+^;*Fat3*^fl/+^ mice. C. VGAT immunostaining for *Isl1*^CRE/+^;*Fat3*^fl/ΔTM^ *Isl1*^cKO^. TdTomato reporter of Cre expression is seen in B’-C’. D. Quantification of the OMPL score for *Isl1*^CRE^ *Fat3*^cKO^. Controls (*Isl1*^CRE/+^;*Fat3*^fl/+^): 0.095 ± 0.065 (n=14 sections, N=3 animals); *Isl1*^cKO^ (*Isl1*^CRE/+^;*Fat3*^fl/ΔTM^): 0.825 ± 0.074 (n=19 sections, N=3 animals), Mann-Whitney test. E. DAPI and Bhlhb5 immunostaining of control retinas (*Isl1*^CRE/+^;*Fat3*^fl/+^). F. DAPI and Bhlhb5 immunostaining of *Isl1*^CRE^ *Fat3*^cKO^ (*Isl1*^CRE/+^;*Fat3*^fl/ΔTM^) retinas. G. Quantification of the number of nuclei per field in the IPL. Controls (*Isl1*^CRE/+^;*Fat3*^fl/+^): 1.25 ± 0.35 (n=12 sections, N=3 animals); *Isl1*^cKO^ (*Isl1*^CRE/+^;*Fat3*^fl/ΔTM^): 5.11 ± 0.39 (n=18 sections, N=3 animals). *t* test. H. Quantification of the number of Bhlhb5+ nuclei per field in the IPL and GCL. Controls (*Isl1*^CRE/+^;*Fat3*^fl/+^): 3.67 ± 0.43 (n=12 sections, N=3 animals); *Isl1*^cKO^ (*Isl1*^CRE/+^;*Fat3*^fl/ΔTM^): 9.33 ± 0.67 (n=18 sections, N=3 animals). *t* test. I. Flicker ERG amplitude at 30 Hz for control (*Isl1*^CRE/+^;*Fat3*^fl/+^, n=6 eyes) and *Isl1*^cKO^ (*Isl1*^CRE/+^;*Fat3*^fl/ΔTM^, n=6 eyes). Unpaired two-tailed Student’s t test. J. Flicker ERG implicit time at 20 Hz for control (*Isl1*^CRE/+^;*Fat3*^fl/+^, n=6 eyes) and *Isl1*^cKO^ (*Isl1*^CRE/+^;*Fat3*^fl/ΔTM^, n=6 eyes). Unpaired two-tailed Student’s t test. K. Representative flicker ERG raw traces for control and *Isl1*^CRE^ *Fat3*cKO. Scale bars: 20µm.

Thus, the high frequency light response defects are not due to the gross disorganization of retinal lamination and do not reflect a role for FAT3 in RGCs or ON-CBCs alone. The 30 Hz amplitude and 20 Hz implicit timing were also unaffected by removal of *Fat3* from GABAergic ACs and the type 2 OFF-CBC subset using *Bhlhe22*^CRE^ (**Figure S3A-K**). We were not able to directly test whether high frequency light response requires FAT3 in OFF-CBCs, as there is no Cre line that is active in all OFF-CBCs. Nonetheless, these experiments showed that the high frequency light response defects are not caused by retinal lamination defects and most likely involve the loss of FAT3 from OFF-CBCs.

To investigate possible cellular origins of the visual deficits, we analyzed cone and BC number and morphology. We found no change in the number or distribution of cones (detected by staining for cone arrestin, ARR3) (27.77 ± 0.91 cones, n=13 sections from 4 eyes vs. 29.00 ± 0.59 cones, n=15 sections from 4 WT eyes) or BCs (detected by staining for VSX2), which occupied a similar area in WT compared to *Fat3*^ΔTM/ΔTM^ mutant retinas (21.09 ± 1.04 arbitrary units, n=15 sections from 4 eyes vs. 21.59 ± 0.97 arbitrary units, n=15 sections from 4 WT eyes (**Figure S4A-F**). To visualize OFF-CBC morphology, we created a novel AAV vector (AAV8-Grik1-GFP) using our previously described *Grik1* enhancer/promoter element^17^. WT and mutant P2/P3 retinas were injected with AAV8-Grik1-GFP and evaluated at P22. Whereas 100% (n=12 cells from 5 animals) of OFF-CBC axons terminated and stratified in the OFF sublaminae of the IPL in WT animals, only 49 ± 17 % of the mutant OFF-CBC axons (n=7 animals, 22 cells) terminated properly with 56 ± 19 % of axons terminating instead in the OMPL or in both the OMPL and the IPL (**Figure S4G-I**). Despite this change in axon position, mutant OFF-CBCs showed typical BC morphologies and extended dendrites that terminated properly in the OPL, as in the WT retina (**Figure S4G-I**). Thus, unlike its role in ACs, FAT3 is not essential for the position of OFF-CBC dendritic arbors.

### FAT3 intracellular signaling is critical for high frequency light response and the step ERG d-wave

FAT3 is a versatile protein that can mediate non-autonomous interactions via its extracellular domain and autonomous effects by recruiting different sets of cytoskeletal effectors to its ICD^6^. To better understand how FAT3 signaling supports high frequency visual signal transmission and contributes to the step ERG d-wave, we analyzed *Fat3*^ΔICD-GFP^ animals, in which most of the ICD is replaced with GFP while keeping the extracellular and transmembrane domains anchored to the cell membrane^6^ (**Figure 5A**).

**Figure 5:**
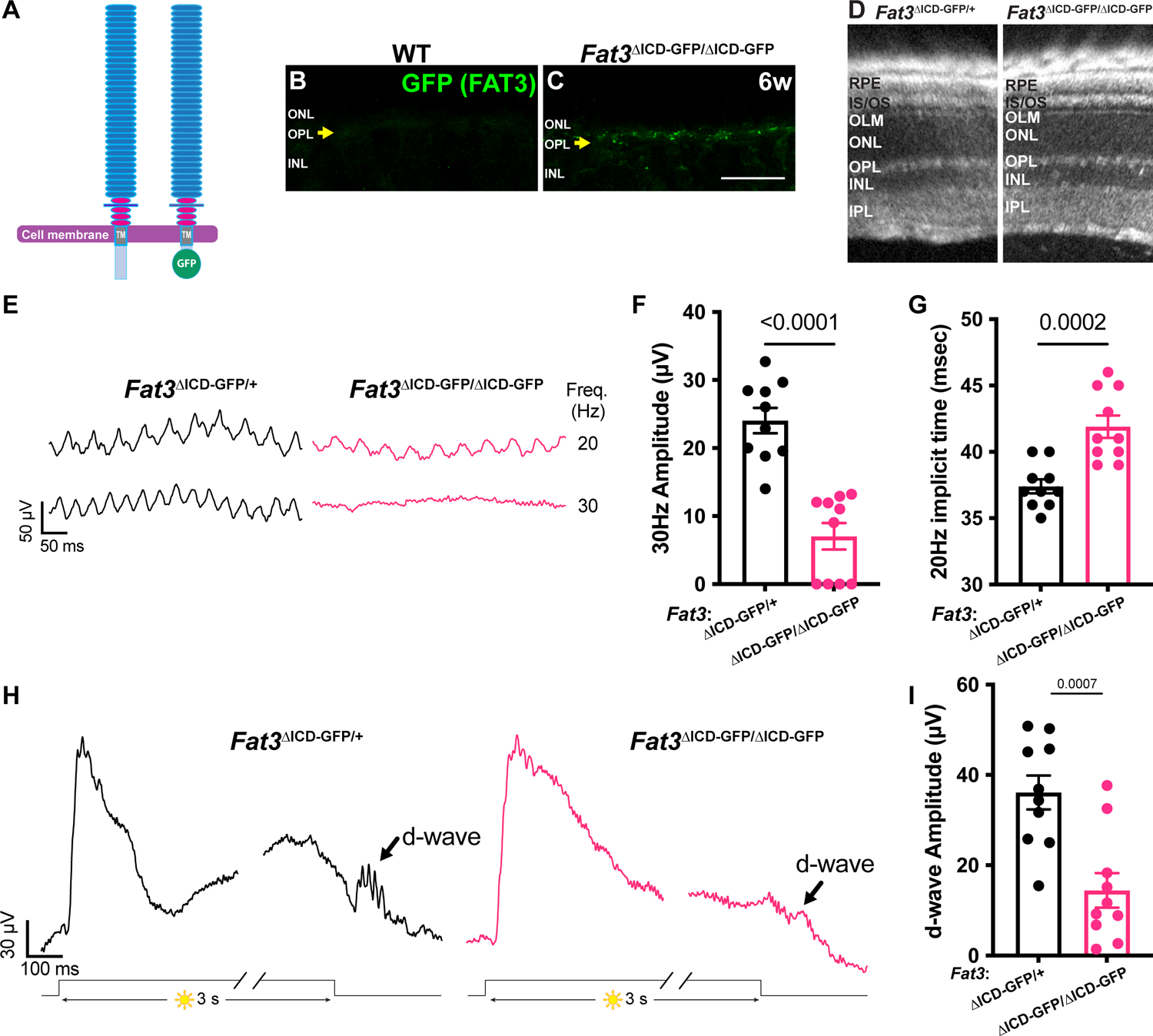
High frequency flicker ERG and step ERG of FAT3 intracellular domain (ICD) deficient mice. A. Schematics of molecular structure of FAT3 wild type protein and FAT3^ΔICD-GFP^. B. Immunostaining for GFP in WT retinal sections. The arrow points the OPL. C. Immunostaining for GFP in *Fat3*^ΔICD-GFP/ΔICD-GFP^ retinal sections. The arrow points the OPL. D. Representative OCT images of *Fat3*^ΔICD-GFP/+^ control and *Fat3*^ΔICD-GFP/ΔICD-GFP^ eyes. E. Representative flicker ERG raw traces of *Fat3*^ΔICD-GFP/+^ control and *Fat3*^ΔICD-GFP/ΔICD-GFP^ eyes elicited by 3.162 cd s/m^2^ flashes at 20 and 30 Hz frequencies. F. Flicker ERG amplitude at 30 Hz for *Fat3*^ΔICD-GFP/+^ control (n=10 eyes) and *Fat3*^ΔICD-GFP/ΔICD-GFP^ (n=10) eyes. Unpaired two-tailed Student’s t test. G. Flicker ERG implicit time at 20 Hz for *Fat3*^ΔICD-GFP/+^ control (n=10 eyes) and *Fat3*^ΔICD-GFP/ΔICD-GFP^ (n=10) eyes at 20 Hz. Unpaired two-tailed Student’s t test. H. Representative step ERG raw traces of *Fat3*^ΔICD/+^ control and *Fat3*^ΔICD-GFP/ΔICD-GFP^ eyes elicited by a 3-second step light at 1000 cd/m^2^ intensity. I. Statistics of step ERG d-wave amplitudes for *Fat3*^ΔICD-GFP/+^ control (n=10 eyes) and *Fat3*^ΔICD-GFP/ΔICD-GFP^ (n=10) eyes elicited by a 3-second step of light at 1000 cd/m^2^ intensity. Unpaired two-tailed Student’s t test.

These animals showed expression of the FAT3-GFP fusion protein in the OPL (**Figure 5B,C**), consistent with FAT3 protein localization (**Figure 3D**). In *Fat3*^ΔICD-GFP/ΔICD-GFP^ mutant mice, ACs migrate to abnormal cell layers and fail to retract their neurites, but do not form ectopic plexiform layers, as shown previously^6^. Therefore, analyzing *Fat3*^ΔICD-GFP/ΔICD-GFP^ mutants can reveal whether high frequency light response depends on intracellular signaling and/or the ability to form synapses.

Consistent with previous histological analysis^6^, no ectopic plexiform layers were detected by OCT *in vivo* imaging of *Fat3*^ΔICD-GFP/ΔICD-GFP^ eyes (**Figure 5D**). Nonetheless, the amplitude of the 30 Hz flicker ERG response was decreased in *Fat3*^ΔICD-GFP/ΔICD-GFP^ mice, as in *Fat3*^ΔTM/ΔTM^ mice (**Figure 5E,F**). Likewise, the 20 Hz implicit time was delayed and the amplitude of the d-wave in step ERG responses was decreased (**Figure 5G-I**). These changes in *Fat3*^ΔICD-GFP/ΔICD-GFP^ mice are highly similar to those in *Fat3*^ΔTM/ΔTM^ mice (**Figure 1D,E,G**). The loss of high frequency flicker ERG responses even in the absence of ectopic plexiform layers further underscores that changes in AC wiring do not contribute to altered visual function in *Fat3* mutant mice. Rather, these results suggest a potential role for FAT3 signaling in OFF-CBCs, in keeping with the results of the genetic analyses described above.

### GRIK1 localization to the ribbon synapse is reduced in *Fat3* mutants

Given the absence of obvious changes in the position or morphology of BC dendrites (**Figure S4**), we hypothesized that the observed changes in retinal function are due instead to FAT3-dependent effects on retinal ribbon synapses. Indeed, in addition to binding to a variety of cytoplasmic effectors important for neuronal migration and neurite retraction, GST-pulldown assays showed that the FAT3 ICD binds to several proteins associated with synaptic function, such as HOMER1^6^ and the LAR family protein, PTPσ, which is encoded by the *Protein tyrosine phosphatase receptor type S* (*Ptprs*) gene (**Figure 6A**). Further, *Drosophila* Fat-like interacts with the related Receptor-type Protein Tyrosine Phosphatase (RPTP) protein dLAR^19^, and PTPσ is required for excitatory synapse formation in hippocampal neurons^20–22^. Since *Ptprs* RNA is detected in both ON- and OFF-BCs^2^ (**Figure S2**), we asked whether cell type-specific synaptic features might be altered in the absence of FAT3. Immunostaining revealed that PTPσ localized to the post-synaptic dendrites of CBCs, overlapping with the postsynaptic glutamate receptor subunit GRIK1 (**Figure 6B, Figure S5A,B,C-G**) and apposing CtBP2+ ribbons in the ARR3+ cone terminals (**Figure 6C**). However, significantly less PTPσ was detected in the OPL of *Fat3*^ΔTM/ΔTM^ mutants (density in the OPL of 4871 ± 463.5, n=3 mutants vs 8682 ± 583.0 in n=4 WT) (**Figure 6D-F**).

**Figure 6:**
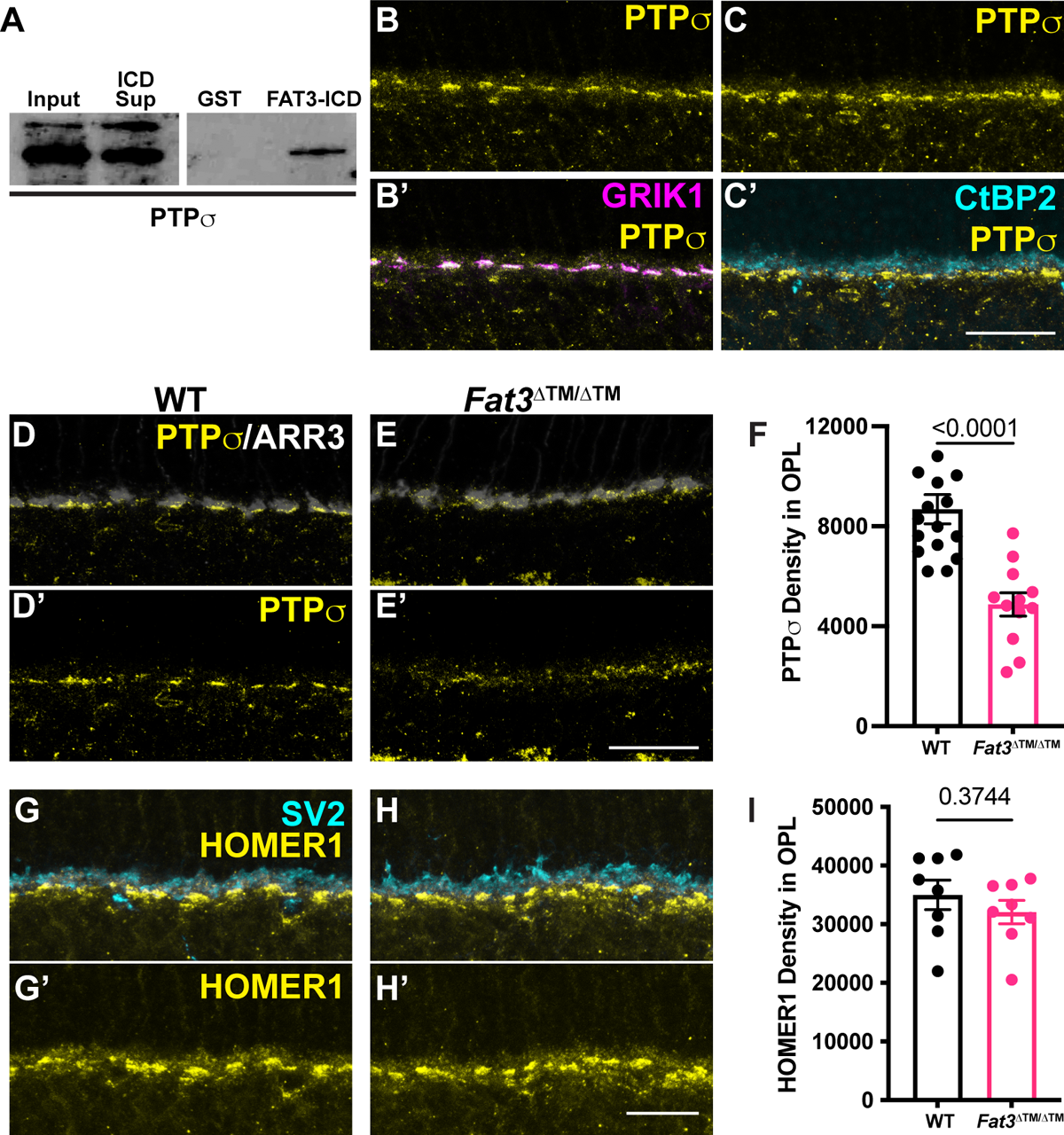
PTP**σ** and HOMER1 localization in WT and FAT3 mutant retinas. A. Binding of FAT3-ICD to PTPσ was assayed using a pull down, with GST fused to FAT3-ICD. B. Immunostaining of PTPσ and (B’) GRIK1. C. Immunostaining of PTPσ and (C’) CtBP2, a marker of ribbons in photoreceptor axons. D. Immunostaining of PTPσ in WT retinas and ARR3 (D’). E. Immunostaining of PTPσ in *Fat3*^ΔTM/ΔTM^ retinas and ARR3 (E’). F. Quantification of PTPσ integrated intensity in the OPL. WT Controls: 8682 ± 583 (n=16 sections, N=4 animals); *Fat3*^ΔTM/ΔTM^: 4871 ± 463.5 (n=12 sections, N=3 animals), Mann-Whitney test. G. Immunostaining of HOMER1 and SV2, and (G’) HOMER1 alone in WT retinas. H. Immunostaining of HOMER1 and SV2, and (H’) HOMER1 alone in *Fat3*^ΔTM/ΔTM^ retinas. I. Quantification of HOMER1 integrated intensity in the OPL. WT Controls: 34997 ± 2509 (n=8 sections, N=3 animals); *Fat3*^ΔTM/ΔTM^: 32051 ± 2003 (n=8 sections, N=3 animals), t test. Scale bars: 20µm.

Immunostaining revealed no change in the levels of post-synaptic HOMER1, which continued to oppose SV2+ synaptic terminals in the OPL of *Fat3* mutants (**Figure 6G-I**). Pre-synaptic CTBP2+ ribbons were also not affected in *Fat3* mutants (OPL mean fluorescence intensity: 5.00 ± 0.14 in n= 4 *Fat3*^ΔTM/ΔTM^ animals *vs.* 4.80 ± 0.16 in n=4 WT; 9.70 ± 0.39 in n= 4 *Fat3*^ΔICD-GFP/ΔICD-GFP^ animals *vs.* 10.03 ± 0.32 in n=4 WT) (**Figure S5H-M**), suggesting that FAT3 may not be required for synapse formation *per se*.

GRIK1 is the only subunit of ionotropic glutamate receptors (iGluR) enriched specifically in OFF-CBCs ^2,17,23,24^ (**Figure S5C,E,F**). GRIK1 staining was strongly reduced in the OPL of both *Fat3*^ΔTM/ΔTM^ (**Figure 7A-C**) and *Fat3*^ΔICD-GFP/ΔICD-GFP^ (**Figure 7D-F**) retinas compared to WT littermates (density in the OPL: 25838 ± 5028, n=4 *Fat3*^ΔTM/ΔTM^ vs 52727 ± 7204 in n=4 WT and 11687 ± 1110 in n= 4 *Fat3*^ΔICD-GFP/ΔICD-GFP^ *vs.* 16119 ± 1168 in n=4 WT). BCs also expressed less *Grik1* RNA in *Fat3*^ΔTM/ΔTM^ animals compared to WT (**Figure S6A-C**). By contrast, the ON-CBC synaptic protein GRM6, which is a metabotropic glutamate receptor subunit (**Figure S5D,G**), was present at similar levels in the OPL of mutant and control retinas (**Figure S6E,F,I**) and in *Fat3*^ΔICD-GFP/ΔICD-GFP^ mutants (**Figure S6G,H,J**). There was also no difference in the level of *Grm6* RNA in the INL of mutant and control retinas (**Figure S6A,B,D**). Loss of *Ptprs* also resulted in a modest decrease of GRIK1 in the OPL (**Figure 7G-I**). However, unlike *Fat3* mutants, *Ptprs*^-/-^ animals did not show any change in the ERG response to a 30 Hz flickering stimulus, the implicit timing of the response to a 20 Hz stimulus, or the d-wave amplitude in the step ERG assay (**Figure S6K-O**). Thus, FAT3 likely impacts not only the presence of PTPσ and GRIK1, but also additional features of the CBC synapses necessary for responses to high temporal frequency stimuli.

**Figure 7:**
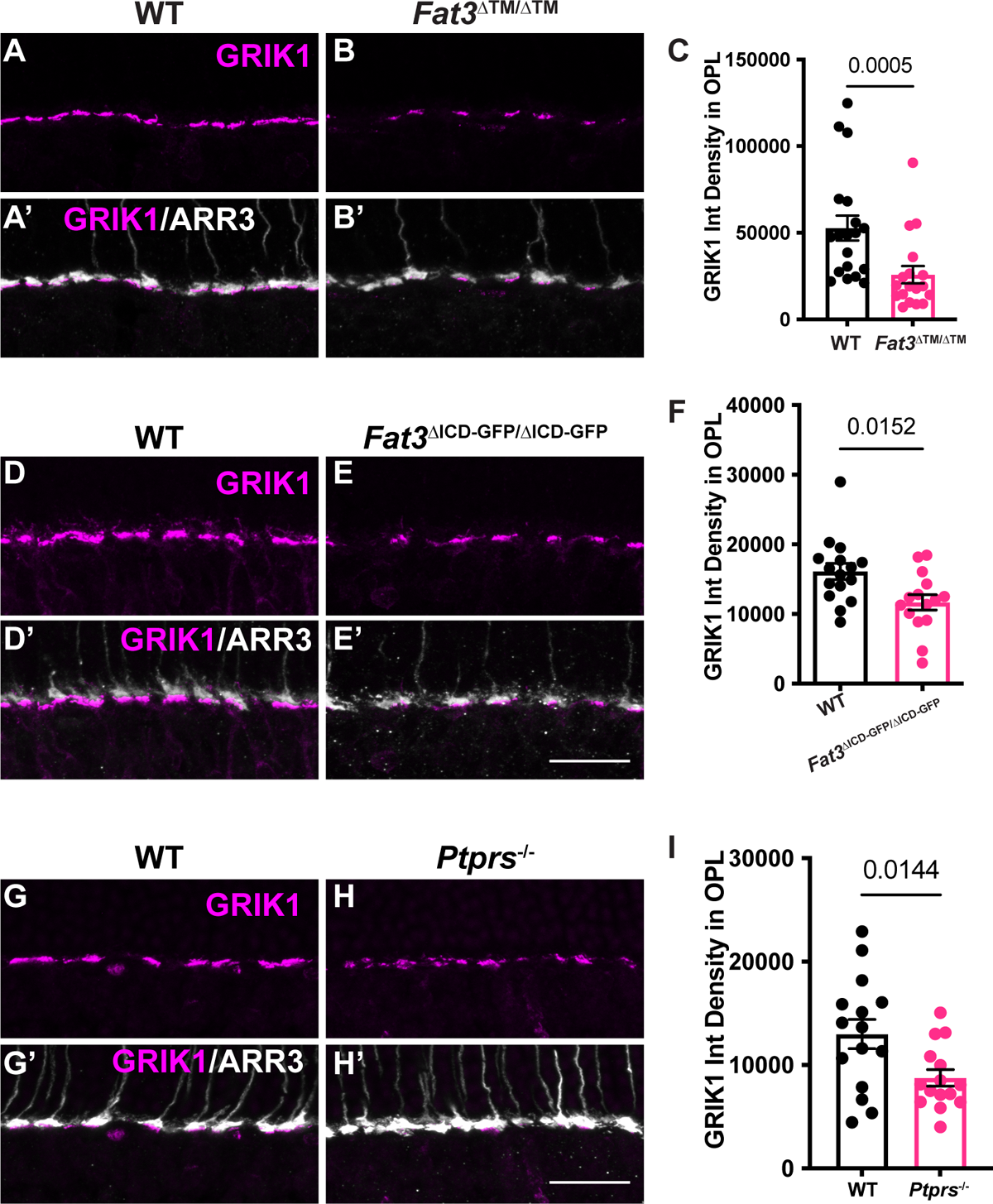
GRIK1 localization in WT, FAT3 and PTP**σ** mutant retinas. A. Immunostaining for GRIK1 in WT retinas. B. Immunostaining for GRIK1 in *Fat3*^ΔTM/ΔTM^ retinas. Cone arrestin (ARR3) labels the cone photoreceptor axonal endings in the OPL in A’ and B’. C. Quantification of GRIK1 integrated intensity in the OPL. WT Controls: 52727 ± 7204 (n=19 sections, N=4 animals); *Fat3*^ΔTM/ΔTM^: 25838 ± 5028 (n=18 sections, N=4 animals), Mann-Whitney test. D. Immunostaining for GRIK1 in WT retinas. E. Immunostaining for GRIK1 in *Fat3*^ΔICD-GFP/ΔICD-GFP^ retinas. Cone arrestin (ARR3) labels the cone photoreceptor endings in the OPL in D’ and E’. F. Quantification of GRIK1 integrated intensity in the OPL. WT Controls: 16119 ± 1168 (n=16 sections, N=4 animals); *Fat3*^ΔICD/ΔICD^: 11687 ± 1110 (n=15 sections, N=4 animals), Mann-Whitney test. G. Immunostaining for GRIK1 in WT retinas. H. Immunostaining for GRIK1 in *Ptprs*^-/-^ retinas. Cone arrestin (ARR3) labels the cone photoreceptor endings in the OPL in G’ and H’. I. Quantification of GRIK1 integrated intensity in the OPL. WT Controls: 12996 ± 1418 (n=15 sections, N=3 animals); *Ptprs*^-/-^: 8758 ± 792 (n=15 sections, N=3 animals), t test. Scale bars: 20µm.

## Discussion

A major goal in retinal physiology is to understand how the diverse cell types of the retina, including >80 types of interneurons, properly connect to form and operate within parallel circuits that transform information about specific elements of the visual world. A valuable approach is to characterize retinal physiology and visual behavior in mice carrying mutations in genes required for proper connectivity and/or function.

However, the nature of the retinal neurons that enable responses to stimuli that change with high temporal frequency has been explored very little, in part because standard assays of retinal function, particularly the conventional scotopic and photopic ERGs, are not designed to detect signals elicited by these stimuli. Despite lacking the cellular level resolution that can be achieved by *ex vivo* patch clamping, the ERG is a relatively simple measurement that can be used *in vivo* to reveal behaviorally-relevant changes in overall retinal function from populations of cells. Here, we use flicker and step ERGs to show that retinal activity in response to high frequency flickering and light-off stimuli is altered in *Fat3* mutant mice, which also show an impaired behavioral response to flickering stimuli. FAT3 is a multi-functional transmembrane protein that is expressed by ACs, BCs, and RGCs and is required for proper lamination of retinal neurons and their synapses. By analyzing the consequences of *Fat3* deletion from different retinal cell types, we discovered that the loss of high frequency light response was not due to changes in AC position or connectivity. Instead, these visual defects appear to be due to abnormal BCs. Although the precise nature of the synaptic defects remains unclear, we found that FAT3 binds to the synaptic protein PTPσ and that both FAT3 and PTPσ are required for enrichment of GRIK1 at the cone to OFF-CBC synapse. Together, these data uncover a role for FAT3 in the formation and/or maintenance of functional cone to BC synapses.

### FAT3 mutant mice show deficits in high frequency light response

Despite their severe defects in neuronal wiring, *Fat3* mutant mice are not blind, as seen by the preservation of conventional ERG and optomotor responses. However, when presented with fast flickering stimuli, *Fat3* mice showed severe visual deficits consistent with loss of signaling from cones to OFF-CBCs. In support of this idea, abnormal high temporal frequency light response was observed only in mice lacking *Fat3* in BCs and did not correlate with effects on retinal lamination (**Figure 1,2,4**). Additionally, *Fat3* is expressed by multiple BC types, including GRIK1*-*positive OFF-CBCs, and GRIK1 levels were reduced in the OPL of *Fat3* mutant mice (**Figure S2,3,7**), as were levels of PTPσ, a synaptic protein that binds to FAT3 (**Figure 6**). To independently assay cone to OFF-CBC signaling, we established a simple and non-invasive step ERG protocol and showed that the d-wave was reduced in *Fat3* mutant mice compared to WT littermates (**Figure 1**). Collectively, these data suggest that FAT3 is required for proper OFF-CBC function and hence the ability to detect fast flickering stimuli. This phenotype is fundamentally different from that which occurs in *Fat3* mutant ACs, which extend extra neurites that form ectopic synapses.

BCs are the first retinal interneurons that encode, segregate, and relay visual information into over a dozen pathways for further processing. It had been hypothesized that the OFF-CBCs mediate high frequency visual signal transmission, based upon indirect evidence from flicker ERG studies of mice carrying mutations such as *Grm6*^-/-9^. OFF-CBC activities are thought to be initiated by two classes of ionotropic GluRs, namely kainate and AMPA receptors, which were proposed to differentially encode temporal signals from cones^25^. AMPA receptors also have been reported to mediate high frequency signaling in cb2, a particular OFF-CBC subtype in ground squirrel retina, as shown by *ex vivo* patch recording^26^. However, due to lack of reliable antibodies, we were not able to investigate *Fat3* mutant mice for changes in AMPA receptor level or localization.

It is unclear whether the observed changes in ERG responses are entirely due to altered OFF-CBC activity. In addition to OFF-CBCs, *Fat3* RNA is weakly expressed in the 5D and 6 ON-CBC subtypes^2^ (**Figure S2**), possibly affecting some ON-CBC functions directly. However, it is unlikely that an autonomous change in only two subtypes would alter the ON-CBC population responses that the ERG measures.

Further, *Fat3* mutants showed no detectable change in the level of GRM6, the glutamate receptor that leads to membrane voltage changes in ON-CBCs^27^. Non-autonomous effects are possible, as OFF-CBCs are connected to ON-BCs via AII amacrine cells in IPL. Thus, modulation originating within the IPL could affect OPL signaling through a backpropagation mechanism, which has not been described but was suggested by an *ex vivo* study using pharmacological blockers within the amphibian retina^28^. For example, PTPσ might serve as an intermediary, as it can act in *trans*, is expressed in both ON and OFF-BCs^2^ (**Figure S2**), and is reduced in *Fat3* mutants (**Figure 6**). Further studies, including detailed patch clamp analysis, are needed to address these possibilities.

### *Fat3* effects on high temporal frequency light response requires intracellular signaling

The discovery of a new type of synaptic defect in *Fat3* mutant retinas highlights FAT3’s versatility as a signaling molecule. Previous work showed that FAT3 acts through different motifs in its ICD to control AC migration, neurite retraction, and synapse localization, likely by recruiting different combinations of cytoplasmic effectors^6^. Here, we found that the FAT3 ICD is also required for proper organization of the synapses between cones and OFF-CBCs, possibly acting through a separate module of synapse-related effectors. The FAT3-ICD interacts with several known synaptic proteins, including PTPσ, which is one of the four type IIA family of receptor-type protein tyrosine phosphatase in the LAR-RPTP subfamily^29^. Interactions between FAT3 and PTPσ appear to regulate the amount of GRIK1 at the synapse, since PTPσ levels were reduced in the OPL of *Fat3* mutants (**Figure 6**) and GRIK1 levels were reduced in the OPL of both *Fat3* and *Ptprs* mutants (**Figure 7**). Although PTPσ plays a well-established role in differentiation of the pre-synaptic component of excitatory synapses in the brain^20,22,29^, our findings point to a role on the post-synaptic side, as suggested previously^21^. Further, the related protein LAR also can be localized to the post-synaptic compartment and is required for proper surface expression and clustering of AMPA receptors^30^. Thus, it is possible that PTPσ acts similarly to control the distribution of GRIK1 in CBCs, either on its own or in collaboration with FAT3. Since *Ptprs* mutants do not show the same visual deficits as *Fat3* mutants (**Figure S6**), other LAR subfamily members may compensate. This might also explain why *Ptprs* mutants have no obvious changes in retinal lamination^31^. Alternatively, other FAT3-dependent proteins may enable sufficient GRIK1 activity for OFF-CBC signaling in *Ptprs* mutants. Although much remains to be learned about the contribution of FAT3-PTPσ interactions to the synapse, this seems to be a conserved relationship, since *Drosophila* Fat-like and LAR interact to ensure collective cell migration^19^, in this case acting in *trans*^32^.

There are several ways that FAT3 might influence synaptic function. One model is that the FAT3 ICD serves as a scaffold for synaptic proteins that secures them to the OFF-CBC dendrites in the OPL and thus directly shapes visual signal transmission. This fits with the fact that FAT3, PTPσ, and GRIK1 are all localized to the OPL, and that several other synaptic proteins, including the WAVE regulatory complex, also interact with the FAT3 ICD^6^. Alternatively, FAT3 may ensure directed trafficking of PTPσ and other proteins to the synapse, echoing its role as a tissue polarity protein and its ability to promote asymmetric localization of cytoskeletal proteins^8^. These possibilities are not mutually exclusive, as FAT3 could control synapse assembly during development and then maintain the synapse in the mature retina. Finally, FAT3 may impact the synapse through effects on gene expression. In flies, the Fat-like ICD is cleaved and binds to a transcriptional co-repressor to influence gene expression^33^. Since the LAR ICD can be internalized and thus inhibit transcription^34^, FAT3 and PTPσ could cooperate to control expression of synaptic genes. This notion is supported by the observation that *Grik1* RNA is modestly reduced in *Fat3* mutant bipolar cells (**Figure S6**).

### Limitations of the study

Although our data provide strong evidence that FAT3 impacts the OFF-CBC signaling needed for high frequency light response, this study does not show definitively what is wrong at the level of the synapse. The ERG is a measurement of cellular activity across a population of cells, i.e. a group of cells from the same class that function similarly, such as OFF-CBC vs. ON-CBC. ERGs are not designed to detect signals carried within distinct microcircuits that perform transformations of particular features of a visual scene. Additionally, due to the lack of Cre lines specific for all BCs, or only OFF-CBCs or ON-CBCs, it is not possible to analyze the consequences of FAT3 loss in these cell types. Patch clamping is necessary to study the responses of specific BC types at single-cell resolution. Finally, due to the extremely large size of FAT3, we were not able to carry out the biochemical analysis needed to determine precisely how FAT3 affects PTPσ, GRIK1 and other uncharacterized candidate effectors^6^. Our discovery that CBC activity is compromised in *Fat3* mutants sets the stage for more detailed analysis in the future.

## Acknowledgements

We thank members of the Goodrich and Cepko labs, A.P. Sampath (UCLA), and Michael Do (Boston Children’s Hospital) for their insightful comments; Barbara Caldarone, Mouse Behavior Core, Harvard Medical School for assistance on behavioral studies; Paula Montero-Llopis, Microscopy Resources on the North Quad, Harvard Medical School for support on microscopic imaging; C. Wright (Vanderbilt U.), M.E. Greenberg (Harvard Medical School), and Christophe Mulle (University of Bordeaux) for sharing mouse strains; Jeannie Chen (University of Southern California), Frans Vinberg (University of Utah), Shinya Sato (University of California Irvine), Henri Leinonen (University of Eastern Finland), Gabriella Niconchuk, Sylvain Lapan and Emma West (Harvard Medical School), Neal Peachey (Cleveland Clinic), Peter Lukasiewicz and Daniel Kerschensteiner (Washington University in St. Louis) for materials, technical support, and advice. NIH grants K99EY030951 (to Y.X. before June 30, 2022), EY030912 (V.J.K), Howard Hughes Medical Institute (to C.L.C.); The Edward R. and Anne G. Lefler Center and a Harvard Brain Initiative Bipolar Disorders grant (LVG); Lingang Laboratory startup fund (to Y.X. after July 20, 2022); and a Leonard and Isabelle Goldenson Fellowship, Alice and Joseph Brooks Fund Fellowship, and ANID Becas Chile Postdoctoral Fellowship (ECA). The authors also acknowledge support from an RPB unrestricted grant to the Department of Ophthalmology, University of California, Irvine.

## Author contributions

Conceptualization, ECA, LVG, CLC, YX; methodology, SKW, SP, SS, LL, VJK; investigation, ECA, YX; writing, ECA, YX, LVG, CLC, VJK; funding acquisition, LVG, CLC, YX; supervision, LVG, CLC, YX.

## Declaration of interests

The authors declare no competing interests.

## Methods

### Animals

The *Fat3*^ΔTM^ mouse line lacks exon 23, which contains the coding region for the transmembrane domain. Since no ICD anchored to the membrane has been detected^6,7^, this allele is expected to act as a full loss of function. The *Fat3*^floxed^ line contains LoxP sites flanking exon 23^7^. The *Fat3*^ΔICD-GFP^ mouse line has a deletion of most of the FAT3-ICD, which is replaced by GFP. This line possesses a full extracellular domain anchored to the cell membrane^6^. Heterozygous mice were used as breeders to obtain wildtype and knockout littermates. *Grik1*^-/-^ mice were obtained from Christophe Mulle (University of Bordeaux, France)^35^. *Grm6*^-/-^ mice (also known as *Grm6^nob3^*), which was characterized by Maddox and colleagues^36^, were purchased from The Jackson Laboratory (ME, Strain #: 016883). *Ptprs* KO mice were made by Michel Tremblay’s laboratory (McGill University)^37^. Transgenic mice expressing Cre recombinase were obtained from the following sources: *Ptf1a*^CRE^ (C. Wright, Vanderbilt U.)^16^; *Islet1*^CRE^ (a.k.a. *Isl1^tm1(cre)Sev^*/J)^38^ (The Jackson Laboratory, ME. Strain #: 024242); and *Bhlhe22*^CRE^ (a.k.a *Bhlhb5*^CRE^)^39^ (M.E. Greenberg, Harvard Medical School). Mice were maintained on a 12 hour/12 hour light/dark cycle at 18–23 °C and 40–60% humidity.

Animals were handled ethically according to protocols approved by the Institutional Animal Care and Use Committee at Harvard Medical School. Genotyping was done using real time PCR (Transnetyx, Cordova, TN).

### Electroretinography (ERG)

Mice were dark adapted overnight before *in vivo* ERG recordings. Animals were anesthetized with 100/10 mg/kg ketamine/xylazine cocktail and placed on a heating pad. Their pupils were dilated with a drop of 1% tropicamide solution (Bausch + Lomb).

Electrodes were applied to the cornea to pick up the electrical signals from the retina. Eyes were kept moist by a drop of phosphate buffered saline (PBS). With an Espion E3 System (Diagonsys LLC), four types of ERG tests were performed: 1) scotopic test; 2) photopic tests with 1, 10, 100 and 1,000 cd s/m^2^ under a 30 cd/m^2^ background light to saturate the rod responses; 3) flicker tests at 0.5, 10, 20, 30, 40 and 50 Hz; and 4) step-light test with a three-second light step of 1,000 cd/m^2^. The scotopic and photopic ERGs were conducted as described previously^40^. The flicker ERG was recorded using 3.162 cd s/m^2^ flashes as adapted from a published protocol^9^. The step ERG was created for this study to probe the d-wave from OFF-bipolar cells.

### Optical Coherence Tomography (OCT)

OCT of mouse eyes was conducted using an OCT2 system (Phoenix Research Labs), as described previously^40^. OCT imaging was performed on the mice *in vivo* immediately after the ERG tests to confirm the previously observed *ex vivo* histological changes. Before OCT imaging, a drop of GONAK 2.5% hypromellose solution (Akorn) was applied to the eye as the immersion medium with the OCT lens.

### Flicker-light cued fear conditioning assay

The fear conditioning test for high temporal frequency vision was created by modifying previously published protocols^13^. The Med Associates (St Albans, VT) system was used for the tests with four LED lights (two green and two yellow) controlled by a computer software (Med PC). Mice were videotaped through Media Recorder Software and the fear response of freezing was analyzed by researchers who were blinded to the genotypes as a surrogate measurement of memory linked with visual input. On Day 1, the mice were brought to the electric shock cage individually to get familiar with the environment and procedure. They were in the cage for 30 minutes under a dim house light in the background and LEDs turned off (**Figure S1**). On Day 2, mice were brought back to the electric shock cage, first exposed to static green/yellow LED lights for two minutes, followed by 30 seconds of 33 Hz LED flicker light (i.e. the cue). Within the last 2 seconds of the cue, a series of 0.7mA shocks were initiated to trigger the fear memory linked with the cue. This static-flicker-shock cycle was repeated twice more. On Day 3, contextual memory (i.e. context test) was measured by placing the mouse back into the conditioning chamber for three minutes (no electric shock was delivered during this session), and the duration of freezing was recorded. Then, cued memory was measured by placing the mouse into an altered context, which was composed of different tactile and olfactory cues. The amount of freezing in the altered context was measured as a baseline (3 minutes, static light) followed by measurement of freezing during presentation of the cued stimulus (3 minutes, 33 Hz flicker light). The videos were analyzed by an examiner blinded to the genotypes to extract the freezing time of all three conditions (i.e. context, static and 33 Hz flicker).

### Optomotor assay

Using an OptoMotry System (CerebralMechanics), the optomotor assay to measure the visual acuity of mice was conducted as described previously^40^. The testing grates were set with 100% contrast and were moved at 1.5 Hz temporal frequency. The visual acuity (i.e. maximal spatial frequency in the unit of cycle/degree) was tested by an examiner, who was blinded to the genotype. During each testing episode, the examiner reported either “yes” or “no” to a computer program until the threshold of acuity was reached. The parameter of each testing episode (i.e. spatial frequency) was determined by the computer program and blinded to the examiner.

### Dissections and immunohistochemistry

Animals of the desired postnatal age (3 or 6 weeks of age, as indicated) were euthanized by CO_2_ inhalation and cervical dislocation. Extraocular tissue, the cornea and the lens were removed from the eyes and the eyecups were further fixed by immersion in 4 % paraformaldehyde (PFA, EMS Cat#15710) for 30 minutes (min) at room temperature or 15 min on ice. After several washes with PBS buffer, the eyes were submerged in 30% sucrose and kept at 4°C for at least 2 hours (h). After sucrose cryoprotection, eyes were incubated in NEG-50 (VWR, Cat#84000-154) overnight at 4°C and embedded by freezing in a liquid nitrogen vapor bath. Retinal slices were obtained by cryosectioning the eyes at 20 µm thickness and mounting on Superfrost® Plus Micro Slide (VWR, Cat#48311-703). The sections were either stained immediately or stored at −80°C.

For regular immunohistochemistry, NEG-50 was removed by short incubation in PBS and then sections were blocked and permeabilized by incubation in 5% Normal Donkey Serum (NDS, Jackson ImmunoResearch Cat#017-000-121) in Sorensońs supplemented with 0.5% Triton-X for 1-2 h at room temperature. Sections were then incubated in primary antibody diluted in blocking buffer overnight at 4°C. After several washes with PBS, sections were incubated with fluorescent secondary antibodies diluted in 5% NDS in Sorensońs buffer supplemented with 0.02% Triton-X for 1.5-2 h at room temperature. After final washes, sections were mounted in DAPI-Fluoromount-G (SouthernBiotech Cat#0100-20). Primary antibodies used for immunohistochemistry were: rabbit anti-ARR3 (Millipore Sigma, Cat#AB15282), goat anti-Bhlhb5 (1:500; Santa Cruz Biotechnology, Cat#sc-6045), mouse anti-CtBP2 (1:2,000, BD Biosciences, Cat#612044), rabbit anti-dsRed (cross-reacts with TdTomato, 1:1,000, Clontech, Cat#632496), mouse anti-FAT3^6,7^ (1:200), chicken anti-GFP (1:500; Aves, Cat#GFP-1020), mouse anti-GRIK1 (GluR5, 1:200, Santa Cruz Biotechnology, Cat#sc-393420)^41^, sheep anti-GRM6 (1:2,000, a gift from Jeannie Chen, USC and originally developed by Kirill Martemyanov Lab^42^), goat anti-PTPσ (1:200; R&D Systems, Cat# AF3430), rabbit anti-VGAT (1:300; SynapticSystems, Cat#131002), and mouse anti-VSX2 (Chx10, 1:100; Santa Cruz Biotechnology, Cat#sc-365519). All secondary antibodies were diluted 1:1,000 and were: Donkey anti-chicken Alexa Fluor® 488, Donkey anti-goat Alexa Fluor® 568, Donkey anti-mouse Alexa Fluor® 488, Donkey anti-mouse Alexa Fluor® 568, Goat anti-mouse Alexa Fluor® 647, Donkey anti-rabbit Alexa Fluor® 488, Donkey anti-rabbit Alexa Fluor® 568, Donkey anti-rabbit Alexa Fluor® 647, and Donkey anti-sheep Alexa Fluor® 568.

To detect FAT3 on retinal sections, we performed Hybridization Chain Reaction Immunohistochemistry (HCR-IHC)^43^, according to the manufacturer’s instructions (Molecular Instruments, CA). In brief, on day 1 sections were treated similarly to regular immunohistochemistry. After overnight incubation of the primary mouse anti-FAT3^7^ (1:200) antibody, sections were rinsed with PBS-0.1% Tween-20 (PBS-T) and incubated with 1 µg/mL of initiator-labeled anti-mouse secondary antibody (Molecular Instruments) for 1 h at room temperature. Slides were rinsed with PBS-T and a final rinse with 5X Saline-Sodium Citrate buffer with 0.1% Tween-20 (SSC-T) and incubated with amplification buffer (Molecular Instruments) for 30 min at room temperature. H1 and h2 fluorescently-labeled hairpins were separately denaturated at 95 °C for 90 s followed by 30 min incubation at room temperature in the dark. A 60 mM hairpin solution mix was prepared by adding snap-cooled h1 and h2 hairpins to amplification buffer and incubated on the slides over night at room temperature in a dark, humidified chamber. After several washes with SSC-T, sections were mounted with DAPI-Fluoromount-G.

### *In situ* hybridization (RNAscope)

Tissue collection was performed similar as to for immunohistochemistry, except using RNAase-free conditions. For RNAscope *in situ* hybridization, we used RNAscope® Fluorescent Multiplex Reagent Kit v2 (ACD, Cat#323120) assay following the manufactureŕs instructions. In brief, retinal sections were post-fixed in 4% PFA for 15 min at room temperature, treated with hydrogen peroxide for 10 min at room temperature and treated with Protease III for 10 min at 40°C before probe incubation. Probes were obtained from ACD (see Key Resource table). Immunohistochemistry was performed after *in situ* hybridization by rinsing the sections in PBS after the final RNAscope wash and permeabilized and blocked again with 5% NDS/0.5% Triton X-100 Sorensońs buffer, followed by the regular immunohistochemistry protocol.

### *In vivo* viral injection

The AAV-Grik1-GFP plasmid was generated by cloning a previously identified Grik1 enhancer (CRM4)^17^ upstream of a simian virus 40 (SV40) intron, Kozak sequence, GFP coding sequence, woodchuck hepatitis virus post-transcriptional regulatory element (WPRE), and polyadenylation sequence. To produce the AAV8-Grik1-GFP vector, HEK293T cells were triple transfected with a mixture of AAV-Grik1-GFP plasmid, adenovirus helper plasmid, and rep2/cap8 packaging plasmid. Viral particles were harvested from the supernatant 72 hours after transfection and purified using an iodixanol gradient as described previously^44^. The titer of AAV8-Grik1-GFP was determined by comparing SYPRO Ruby (Molecular Probes) staining for viral capsid proteins (VP1, VP2, and VP3) to that of a reference vector with known titer.

To deliver AAV8-Grik1-GFP into the developing retina, we injected into the subretinal space as described previously^45,46^. In brief, neonatal P2-3 mouse pups were anesthetized by chilling on ice. We injected 2.5 × 10^9^ vector genomes (vg) per eye, which is at a titer not toxic to the eye, diluted in PBS and 0.1% Fast Green (for visualization) using a pulled borosilicate glass needle with an opening of 0.5-1mm diameter connected to an Eppendorf FemtoJet injector into the subretinal space. The pups recovered on a warm pad and upon regaining consciousness they were returned to their mother. We then let them develop until performing histological procedures at P22.

### Image acquisition

After immunohistochemistry or RNAscope, retinal sections were imaged within 300 µm from the optic nerve head on a Leica SP8 or a Zeiss LSM800 confocal microscope. The entire sections were imaged in consecutive z-slices separated by 1 µm using a 40x or 63x oil objective. The z stacks were then projected at maximum fluorescence intensity using Fiji/ImageJ.

### Histochemical quantifications

We assigned random numbers to each image to ensure blinded quantifications. Only after the quantification was done, the identity of the images was revealed to assign the values to their corresponding genotype. All the procedures were done under the same technical parameters, and the comparisons were made between control and experimental conditions within the same experiment to avoid batch effects. The animal (N) and sample (i.e. sections, n) numbers, statistical test performed, and *p* values are indicated in figure legends and/or figures.

We assessed AC migration and the “ectopic synapse score” or “OMPL score’ as described previously^6^. To quantify expression of PTPσ, GRIK1 or GRM6 proteins in the OPL, the images were thresholded until background signal in the OPL was not observed. All images from the same experiment were treated the same way using Fiji (ImageJ). Then, the integrated density was measured to quantify protein expression in the same total area of each image on the OPL region (44.69 µm x 17.78 µm). To quantify CtBP2 fluorescence intensity on immunohistochemistry samples we used Fiji (ImageJ) to measure the Mean Gray Value on areas of the OPL by tracing a rectangle that took up most of the OPL height. In addition, the Mean Gray Value was measured on a rectangle traced on the ONL (region where photoreceptors reside and is used as background signal) to normalize the value of the OPL.

### Statistics

To determine significant differences between control and experimental groups, we used Prism6 software for statistical analysis. After applying a D’Agostino-Pearson omnibus normality test to determine Gaussian distribution of the samples, we either used two-tailed *t* test (if the samples followed a Gaussian distribution) or Mann-Whitney test (if the samples did not follow a Gaussian distribution) to calculate the *p* values. For ERG data of more than two groups, one-way ANOVA with Dunnett multiple comparison test was used.

### GST Pull down and Western blot

For binding analysis, we performed Western blots of supernatants after pulling down binding partners from mouse brain protein homogenates with the FAT3-ICD fused to Glutathione-S-transferase (GST) and GST alone generated previously^6^. Samples were denatured at 95°C for 10 mins and subjected to SDS-PAGE in a 4–12% Criterion™ XT Bis-Tris Protein Gel (Bio-Rad) using XT MES Running Buffer (Bio-Rad). After 2 h at 150 V of electrophoresis, the proteins were transferred to Immobilon-P PVSF (0.45µm, Sigma-Millipore) in Tris-Glycine buffer supplemented with 20% methanol for 1 h at 75V. The Immobilon-P membranes were blocked with 5% skim milk in TBS buffer and then incubated with primary antibodies at 4°C overnight. The primary antibody used for Western blots was mouse anti-PTPσ (1:1,000, Medimabs, Cat# MM-0020-P). After several washes with TBS supplemented with 0.5% Tween 20 (Sigma-Aldrich), the membranes were incubated with a secondary goat anti-mouse HRP antibody (Biorad, Cat# 170-6516) diluted 1:2,000 for 1-2 h at room temperature. The signal was developed using Clarity ECL substrate following the manufactureŕs instructions (Bio-Rad). Western blots were done at least twice with similar results.

### External gene expression profile datasets

Gene expression in different types of retinal bipolar cells was analyzed by using the single-cell RNAseq database (https://singlecell.broadinstitute.org/single_cell/study/SCP3/retinal-bipolar-neuron-drop-seq#study-visualize) and a modified R script that were published previously^2^.

## Resource and data availability

The source data for graphs are provided with this paper. For all quantifications, the raw data are shown along with means and standard errors of the means as well as the statistical analyses utilized. The original images used to generate these data are available from the corresponding author upon request. All unique materials generated in this study are available upon request.

## KEY RESOURCES TABLE

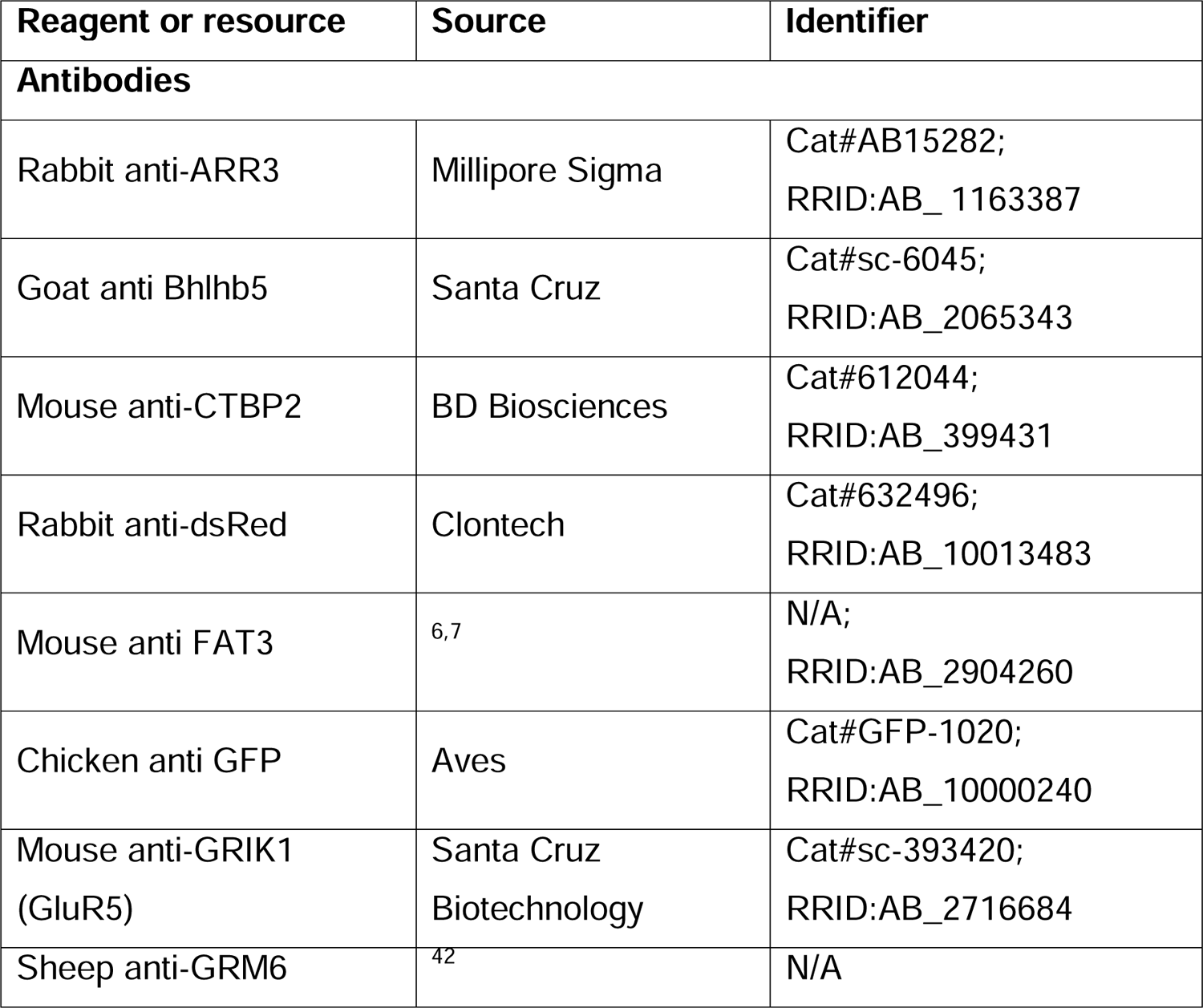
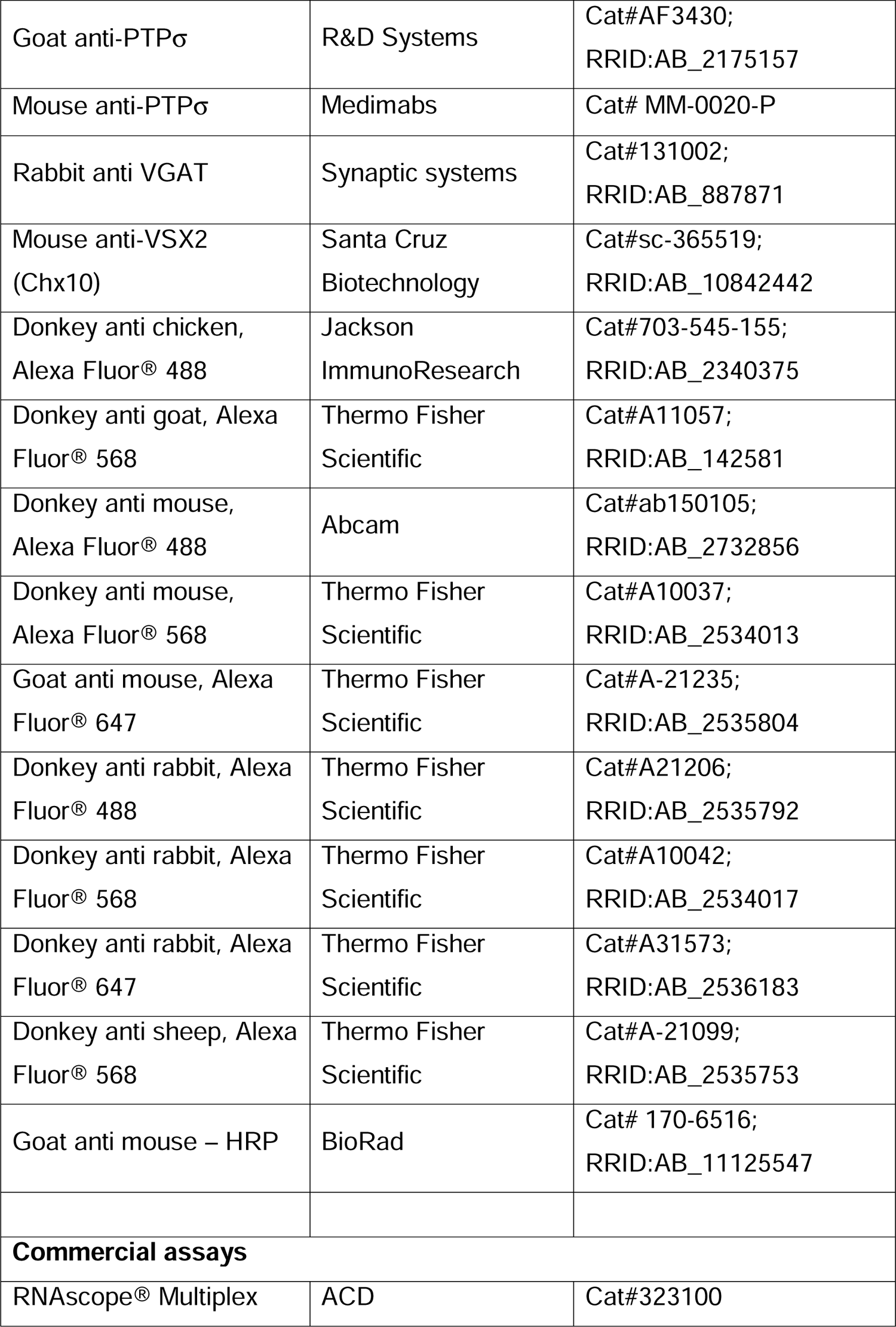
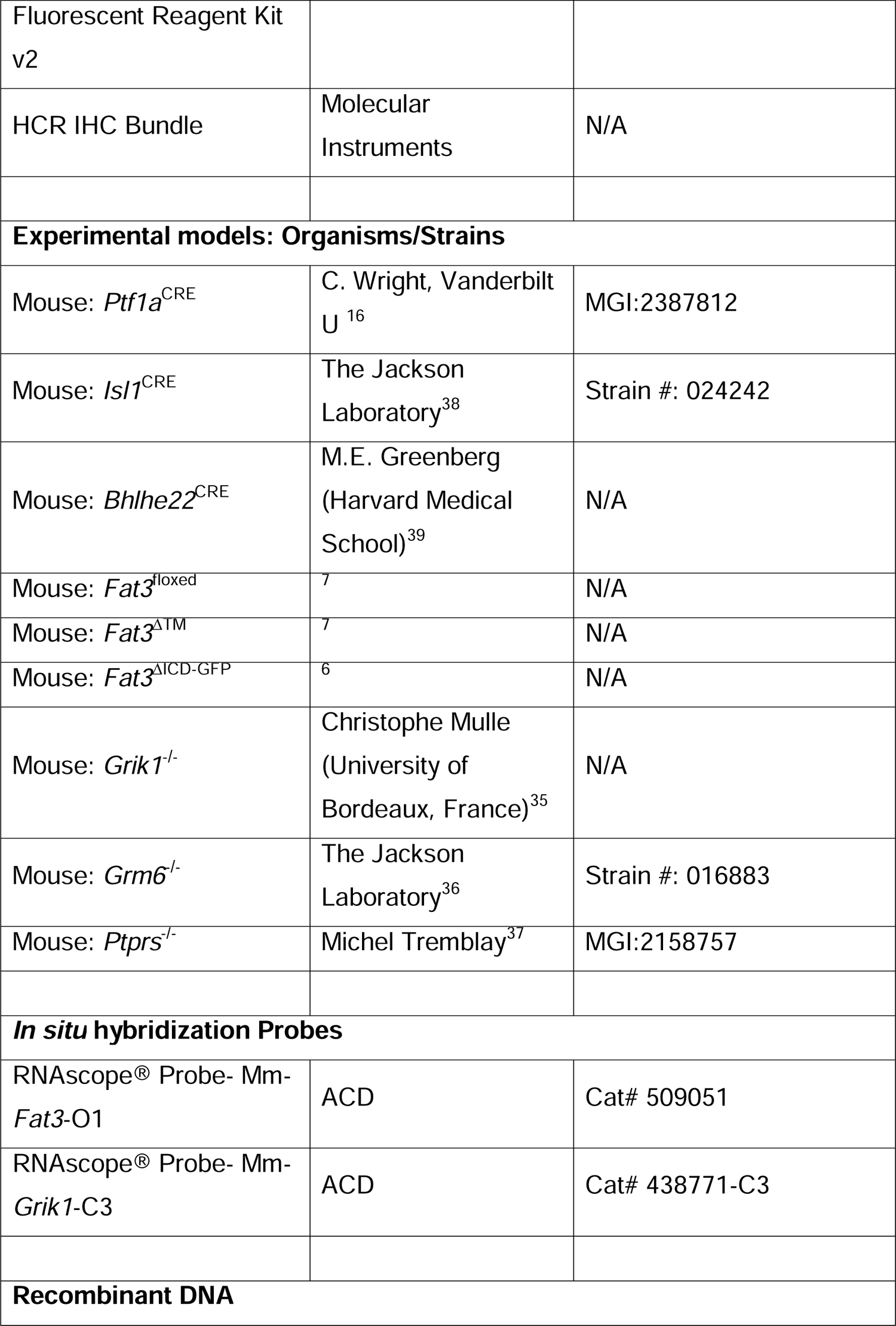
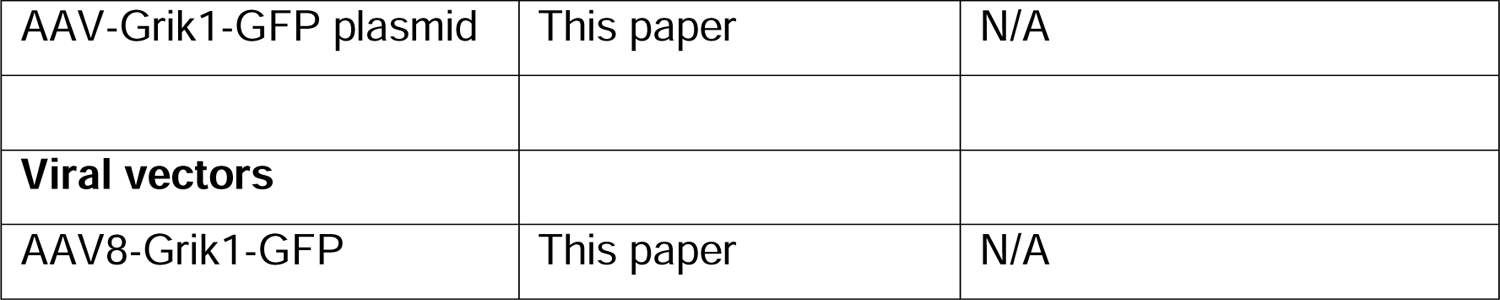

---

**Figure S1** <u>(related to</u> Figure 1): Scotopic and photopic ERG of FAT3-deficient mice. A. Representative OCT images of *Fat3*^ΔTM/+^ control and *Fat3*^ΔTM/ΔTM^ eyes. B. Representative scotopic ERG raw traces of *Fat3*^ΔTM/+^ and *Fat3*^ΔTM/ΔTM^ eyes elicited by 0.1 cd s/m^2^ flashes. C. Statistics of scotopic ERG parameters (amplitude and implicit time of a-wave and b-wave) of *Fat3*^ΔTM/+^ (n=8) and *Fat3*^ΔTM/ΔTM^ (n=10) eyes. D. Representative photopic ERG raw traces of *Fat3*^+/-^ and *Fat3*^-/-^ eyes elicited by 1, 10, 100 and 1,000 cd s/m^2^ flashes at 30 cd/m^2^ background light to saturate the response from rod-pathway. E. Ensemble-averaged photopic ERG b-wave amplitude from *Fat3*^ΔTM/+^ (n=8) and *Fat3*^ΔTM/ΔTM^ (n=10) eyes. F. Schematics of fear conditioning and optomotor behavioral experiment. On Day 1, a mouse is brought to the electric-shock cage with a floor of metal bars for habituation of the environment. On Day 2, the mouse is conditioned by electrical shock paired with 33 Hz flashing light. On Day 3 (see Supplementary movies for representative recordings from Fat3 mutant mice), the mouse is first subjected to a contextual check, in which the “Context” measures the freezing time of the mouse after it is brought back to the electric shock cage, which presents a fear-associated context environment, without the shock. “Static” measures the freezing time of the mouse with a static light, after the covering the metal bars and an odor change. Following this measurement, a 33 Hz flickering light is turned on, and the freezing time of the mouse is measured, as the “flicker” time. G. Fear conditioning responses as freezing time (sec) from *Fat3*^ΔTM/+^ (n=8) and *Fat3*^ΔTM/ΔTM^ (n=9) mice. One-way ANOVA with Dunnett multiple comparison test. H. The visual threshold of spatial frequency of WT (n=8) and *Fat3*^ΔTM/ΔTM^ (n=10) mice measured with the optomotor behavioral assay. Unpaired two-tailed Student’s t test. Abbreviations: RPE, retinal pigmented epithelium; IS/OS, inner-outer segments junction; OLM, outer limiting membrane; OPL, outer plexiform layer; IPL, inner plexiform layer; IMPL: inner misplaced plexiform layer. NS: non-significant. All data are presented as mean ± SEM.

**Figure S2 <u>(related to</u>** Figure 2**): scRNAseq data display.** Expression of *Fat3*, *Grik1*, *Grm6* and *Ptprs* gene transcripts by bipolar cell identity are displayed. These data is based on Shekhar et al. (2016) dataset.

**Figure S3** <u>(related to</u> Figure 4): Effect of removal of *Fat3* from OFF-CBC type 2 on flicker ERG. A. Schematic representation of cells types, i.e. type 2 OFF-CBCs and GABAergic ACs, that lose *Fat3* expression in a *Bhlhe22*^CRE^ cKO. Magenta coloring represents cell type expression of FAT3. B. VGAT immunostaining for control *Bhlhe22*^CRE/+^;*Fat3*^fl/+^ mice. C. VGAT immunostaining for *Bhlhe22*^CRE/+^;*Fat3*^fl/ΔTM^ cKOs. D. Quantification of the OMPL score for *Bhlhe22*^CRE^ *Fat3*cKO. Controls (*Bhlhe22*^CRE/+^;*Fat3*^fl/+^): 0.0 ± 0.0 (n=15 sections, N=3 animals); *Bhlhe22*^cKO^ (*Bhlhe22*^CRE/+^;*Fat3*^fl/ΔTM^): 1.0 ± 0.0 (n=18 sections, N=3 animals), Mann-Whitney test. E. DAPI and Bhlhb5 (a.k.a. *Bhlhe22* gene encoded protein) staining of *Bhlhe22*^CRE/+^;*Fat3*^fl/+^ mouse retinas. F. DAPI and Bhlhb5 staining of *Bhlhe22*^CRE/+^;*Fat3*^fl/ΔTM^ retinas. G. Quantification of the number of nuclei per field in the IPL. Controls (*Bhlhe22*^CRE/+^;*Fat3*^fl/+^): 2.79 ± 0.66 (n=14 sections, N=3 animals); *Bhlhe22*^cKO^ (*Bhlhe22*^CRE/+^;*Fat3*^fl/ΔTM^): 7.53 ± 1.20 (n=15 sections, N=3 animals). Mann-Whitney test. H. Quantification of the number of Bhlhb5+ nuclei per field in the IPL and GCL. Controls (*Bhlhe22*^CRE/+^;*Fat3*^fl/+^): 4.29 ± 0.98 (n=14 sections, N=3 animals); *Bhlhe22*^cKO^ (*Bhlhe22*^CRE/+^;*Fat3*^fl/ΔTM^): 11.13 ± 1.34 (n=15 sections, N=3 animals). *t* test. I. Flicker ERG amplitude at 30 Hz for controls (*Bhlhe22*^CRE/+^;*Fat3*^fl/+^, n=6 eyes) and *Bhlhe22*^CRE^ *Fat3*^cKO^ (*Bhlhe22*^CRE/+^;*Fat3*^fl/ΔTM^, n=6 eyes). J. Flicker ERG implicit time at 20 Hz for controls (*Bhlhe22*^CRE/+^;*Fat3*^fl/+^, n=6 eyes) and *Bhlhe22*^CRE^ *Fat3*^cKO^ (*Bhlhe22*^CRE/+^;*Fat3*^fl/ΔTM^, n=6 eyes). K. Representative flicker ERG raw traces for controls (*Bhlhe22*^CRE/+^;*Fat3*^fl/+^, n=6 eyes) and *Bhlhe22*^CRE^ *Fat3*^cKO^ (*Bhlhe22*^CRE/+^;*Fat3*^fl/ΔTM^, n=6 eyes) at 20 and 30 Hz. Scale bars: 20µm.

**Figure S4** <u>(related to</u> Figure 3<u>, 4)</u>: BC and cones numbers, visualization of *Grik1*+ BCs morphology via injection of an AVV-Grik1-GFP virus in *Fat3* mutants. A. VSX2 immunostaining of WT retinas. B. VSX2 immunostaining of *Fat3*^ΔTM/ΔTM^ retinas. C. ARR3 immunohistochemistry of WT retinas. C’ shows ARR3 staining together with DAPI staining. D. ARR3 immunohistochemistry of *Fat3*^ΔTM/ΔTM^ retinas. D’ shows ARR3 staining together with DAPI staining. E. Quantification of thickness of area occupied by VSX2 staining, as shown in A and B. WT controls: 21.09 ± 1.045 (n=15 sections, N=4 animals); *Fat3*^ΔTM/ΔTM^: 21.59 ± 0.969 (n=15 sections, N=4 animals). t test. F. Quantification of number of cones marked by ARR3, as shown in C and D. WT controls: 27.77 ± 0.907 (n=13 sections, N=4 animals); *Fat3*^ΔTM/ΔTM^: 29.00 ± 0.593 (n=14 sections, N=4 animals). t test. G. Examples of wild type (WT) retinal sections showing cells expressing GFP under the control of the *Grik1* enhancer. The GFP reporter was introduced through *in vivo* injection of AAV8-Grik1-GFP. H. *Fat3*^ΔTM/ΔTM^ retinas injected with AAV8-Grik1-GFP. The OPL is indicated with a yellow arrowhead and the OMPL, labeled with VGAT staining, is marked with a yellow arrow. Scale bar: 20µm. I. Quantification of *Grik1*+ BC dendrites in the OPL (WT Controls: 100%; *Fat3*^ΔTM/ΔTM^: 100%), axons in the IPL (WT Controls: 100%; *Fat3*^ΔTM/ΔTM^: 49 ± 17.4%), and axons in the OMPL (WT Controls: 0%; *Fat3*^ΔTM/ΔTM^: 56 ± 18.5%). Controls: n=5 animals, 12 cells, *Fat3*^ΔTM/ΔTM^: n=7 animals, 22 cells. Mann-Whitney test.

**Figure S5** <u>(related to</u> Figure 6 and 7): Expression pattern of postsynaptic components PTP**σ**, HOMER1, GRIK1 and GRM6 in the retina. A. PTPσ immunostaining in WT retinas B. PTPσ immunostaining in *Ptprs*^-/-^ retinas. Cone arrestin (ARR3) labels the cone photoreceptors endings in the OPL in A’ and B’. C. Schematic representation of OFF-BCs and their synapses with cone photoreceptors. ARR3 (white) labels cones and GRIK1 (magenta) labels postsynaptic BC dendrites. D. Schematic representation of ON-BCs and their synapses with cone photoreceptors. ARR3 (white) labels cones and GRM6 (magenta) labels postsynaptic cone and rod BC (RBC) dendrites. E. GRIK1 and GRM6 (E’) immunostaining of adult WT retina. F. GRIK1 and GRM6 (F’) immunostaining of *Grik1*^-/-^ retina. G. GRIK1 and GRM6 (G’) immunostaining of *Grm6*^-/-^ retina. H. Immunostaining for CtBP2, a marker of presynaptic ribbon in WT retinas. I. Immunostaining for CtBP2 *Fat3*^ΔTM/ΔTM^ retinas. Cone arrestin (ARR3) labels the cone photoreceptors endings in the OPL in H’ and I’. J. Quantification of CtBP2 mean fluorescence intensity in the OPL, normalized by ONL signal. Wild type (WT) Controls: 4.8 ± 0.16 (n=16 sections, N=4 animals); *Fat3*^ΔTM/ΔTM^: 5.0 ± 0.14 (n=16 sections, N=4 animals), *t* test. K. Immunostaining for CtBP2 in WT retinas, littermates of *Fat3*^ΔICD-GFP/ΔICD-GFP^ animals. L. Immunostaining for CtBP2 in *Fat3*^ΔICD-GFP/ΔICD-GFP^ retinas. Cone arrestin (ARR3) labels the cone photoreceptors endings in the OPL in K’ and L’. M. Quantification of CtBP2 mean fluorescence intensity in the OPL, normalized by ONL signal. Wild type (WT) Controls: 10.03 ± 0.32 (n=16 sections, N=4 animals); *Fat3*^ΔICD/ΔICD^: 9.7 ± 0.39 (n=16 sections, N=4 animals), *t* test. Scale bars: 20µm.

**Figure S6** <u>(related to</u> Figure 7): Immunostaining of GRM6 and *in situ* hybridization of *Grik1* and *Grm6* upon loss of *Fat3* in mouse retina. A. VSX2 immunostaining (A) after *in situ* hybridization for *Grik1* (A’) and *Grm6* (A’’) in WT retina. B. VSX2 immunostaining (B) after *in situ* hybridization for *Grik1* (B’) and *Grm6* (B’’) in *Fat3*^ΔTM/ΔTM^ retina. C. Quantification of *Grik1* RNA pixel density in bipolar cells. WT: 352076 ± 27459, n=16 sections, N= 4 animals; *Fat3*^ΔTM/ΔTM^: 264572 ± 18120, n=17 sections, N=4 animals. t test. D. Quantification of *Grm6* RNA pixel density in bipolar cells. WT: 447054 ± 21371, n=16 sections, N= 4 animals; *Fat3*^ΔTM/ΔTM^: 443575 ± 25398, n=17 sections, N=4 animals. t test. E. Immunostaining for GRM6 in WT retinas. F. Immunostaining for GRM6 in *Fat3*^ΔTM/ΔTM^ retinas. Cone arrestin (ARR3) labels the cone photoreceptors endings in the OPL in E’ and F’. G. Immunostaining for GRM6 in WT retinas, littermates of *Fat3*^ΔICD-GFP/ΔICD-GFP^ animals. H. Immunostaining for GRM6 in *Fat3*^ΔICD-GFP/ΔICD-GFP^ retinas. Cone arrestin (ARR3) labels the cone photoreceptors endings in the OPL in H’ and I’. I. Quantification of GRM6 integrated intensity in the OPL. WT Controls: 16435 ± 1953 (n=16 sections, N=4 animals); *Fat3*^ΔTM/ΔTM^: 14603 ± 1195 (n=23 sections, N=5 animals), Mann-Whitney test. J. Quantification of GRM6 integrated intensity in the OPL. WT Controls: 23190 ± 2018 (n=21 sections, N=5 animals); *Fat3*^ΔICD/ΔICD^: 20051 ± 1696 (n=22 sections, N=5 animals), Mann-Whitney test. K. Flicker ERG amplitude at 30 Hz for WT (n=6 eyes) and *Ptprs*^-/-^ retinas (n=6 eyes). L. Flicker ERG implicit time at 20 Hz for WT (n=6 eyes) and *Ptprs*^-/-^ retinas (n=6 eyes). M. Representative flicker ERG raw traces of wild type control and *Ptprs*^-/-^ eyes elicited by 3.162 cd s/m^2^ flashes at 20 and 30 Hz frequencies. N. Statistics of step ERG d-wave amplitudes of WT and *Ptprs*^-/-^ eyes elicited by a 3-second step of light at 1000 cd/m^2^ intensity. O. Representative step ERG raw traces of wild type control and *Ptprs*^-/-^ eyes elicited by a 3-second step light at 1000 cd/m^2^ intensity. Scale bars: 20µm.

## Notes

### Competing Interest Statement

The authors have declared no competing interest.

### Summary of Updates

ERG results of Grik1 and Grm6-KO mice are remove. Some of other data are re-organized.

