## Supplementary figures and images for "High temporal frequency light response in mouse retina requires FAT3 signaling in bipolar cells"

### Supplemental Figures S1-6

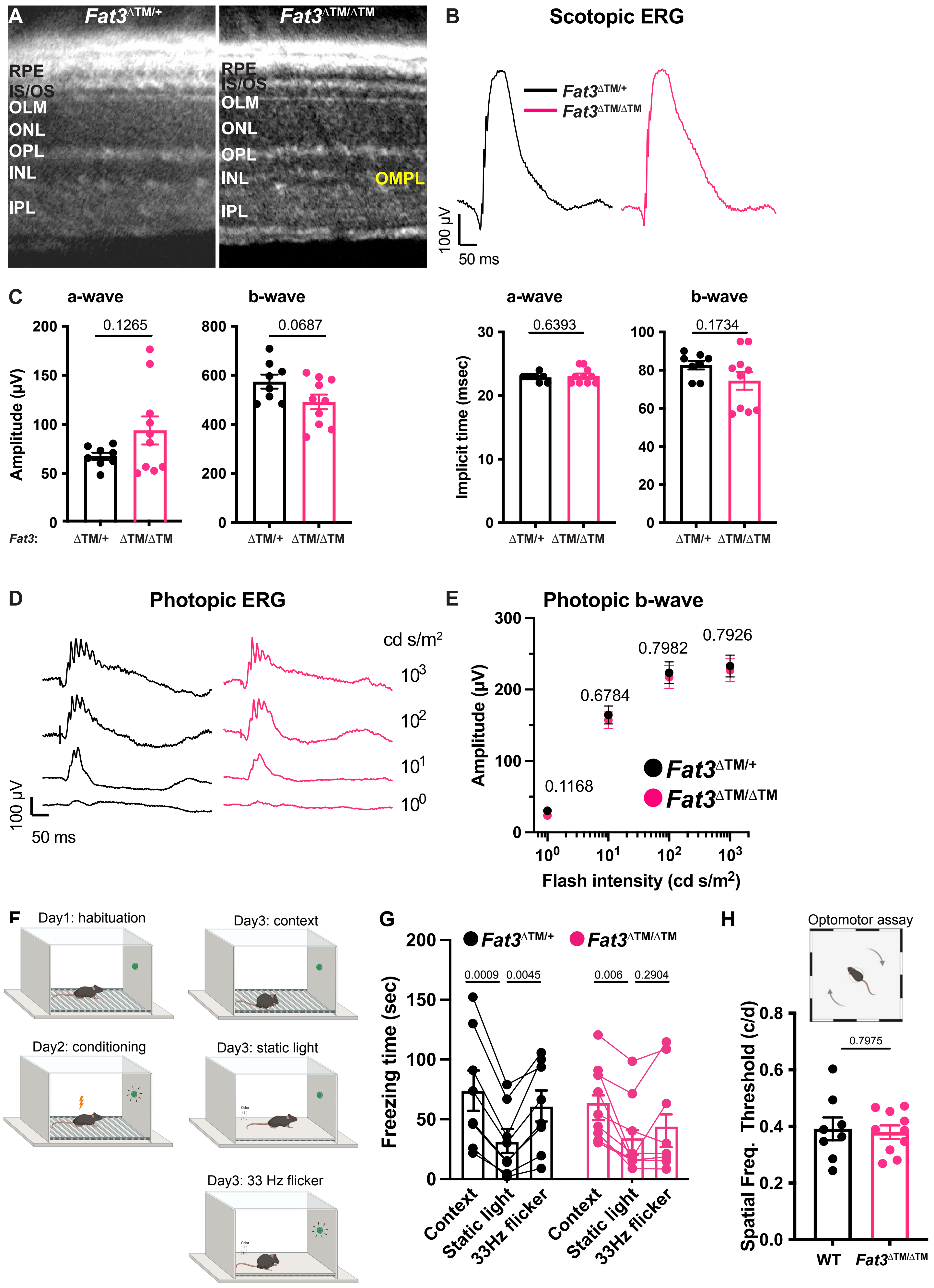

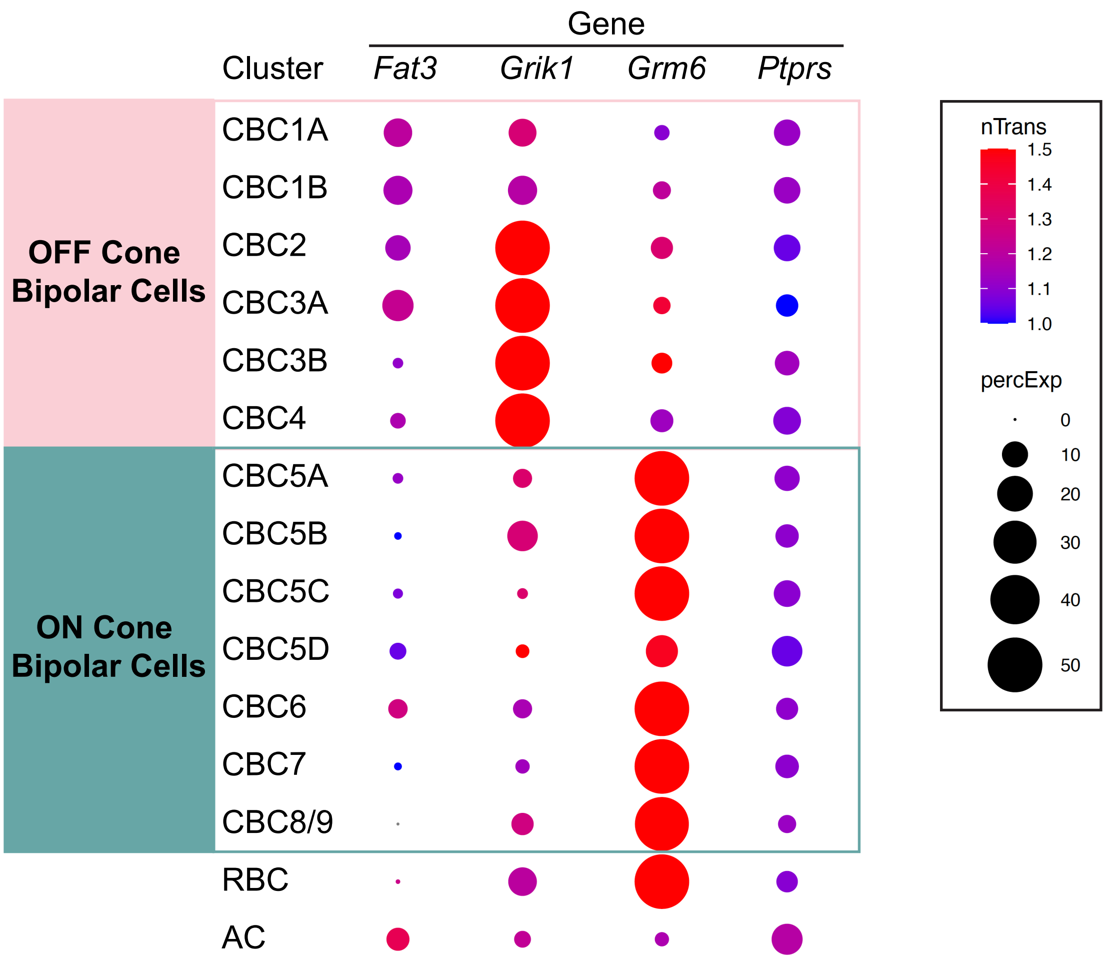

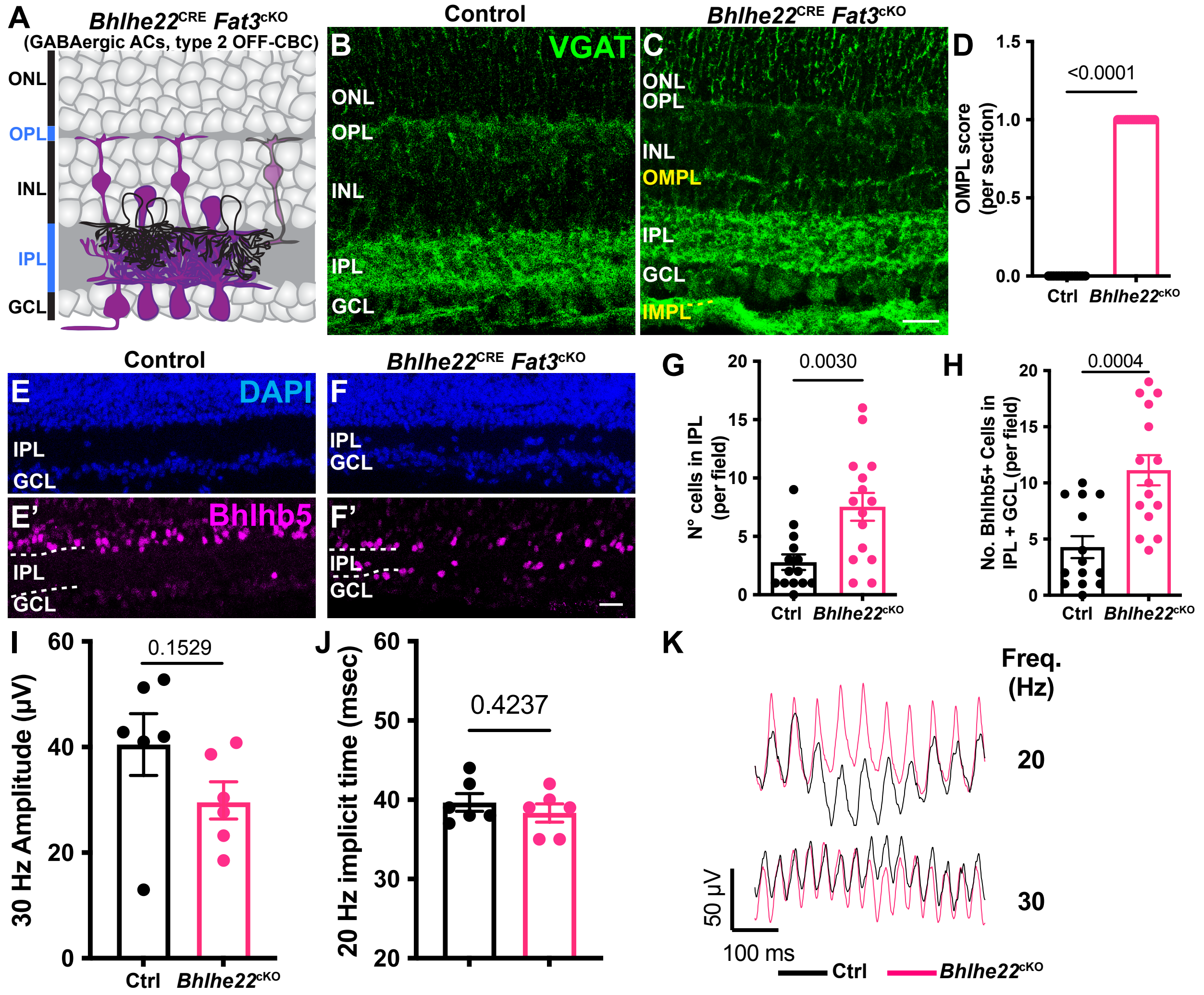

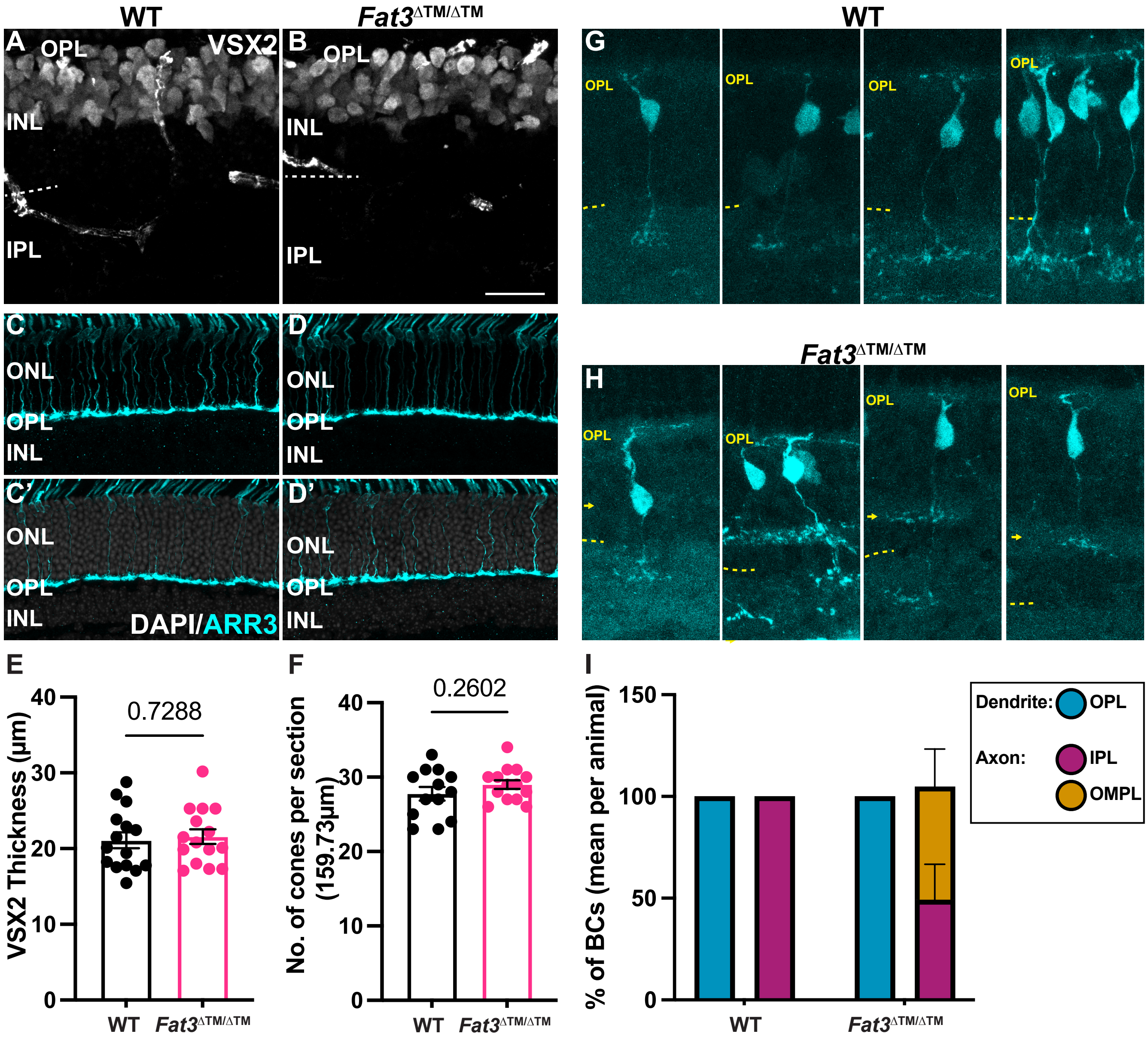

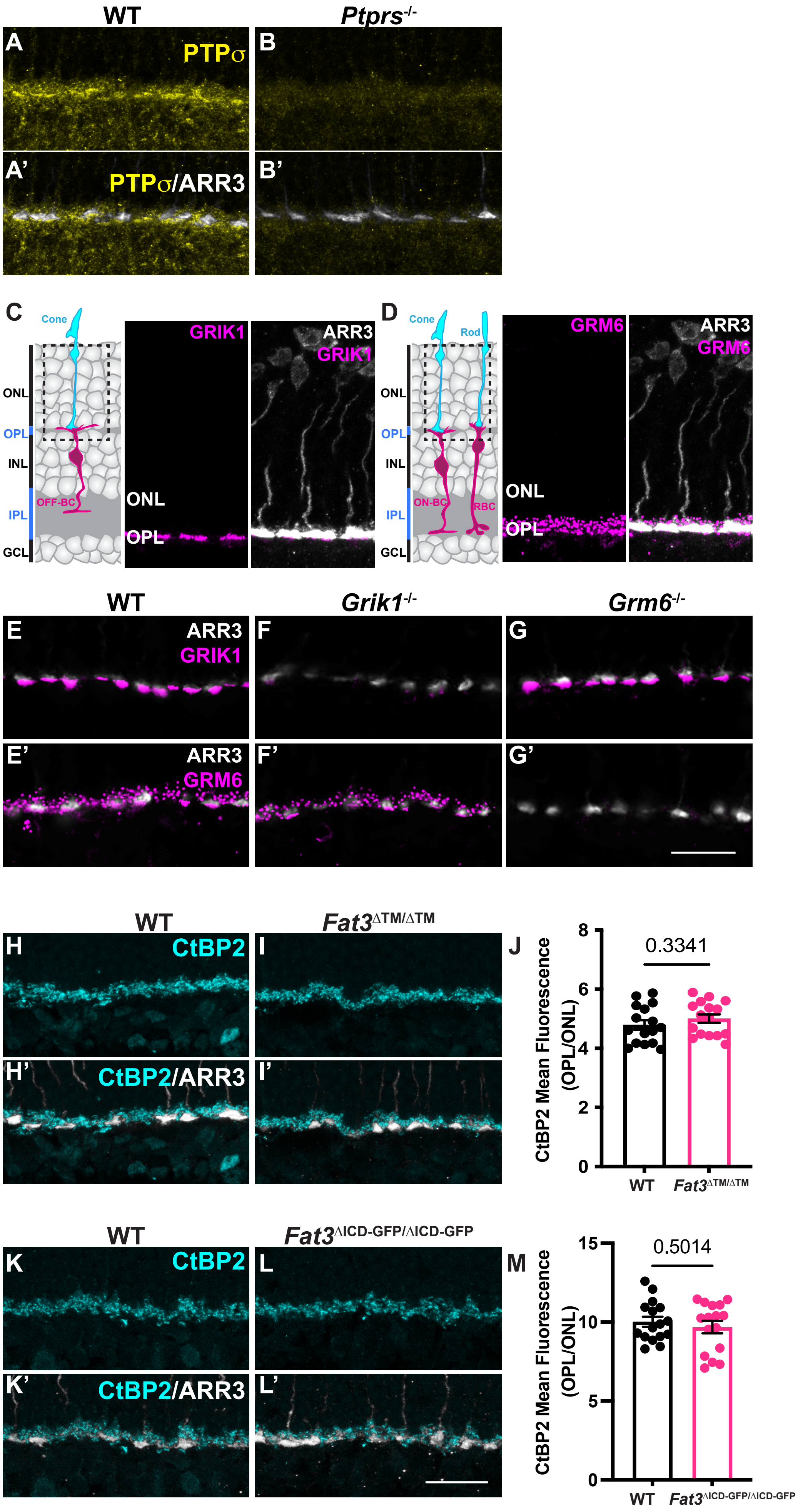

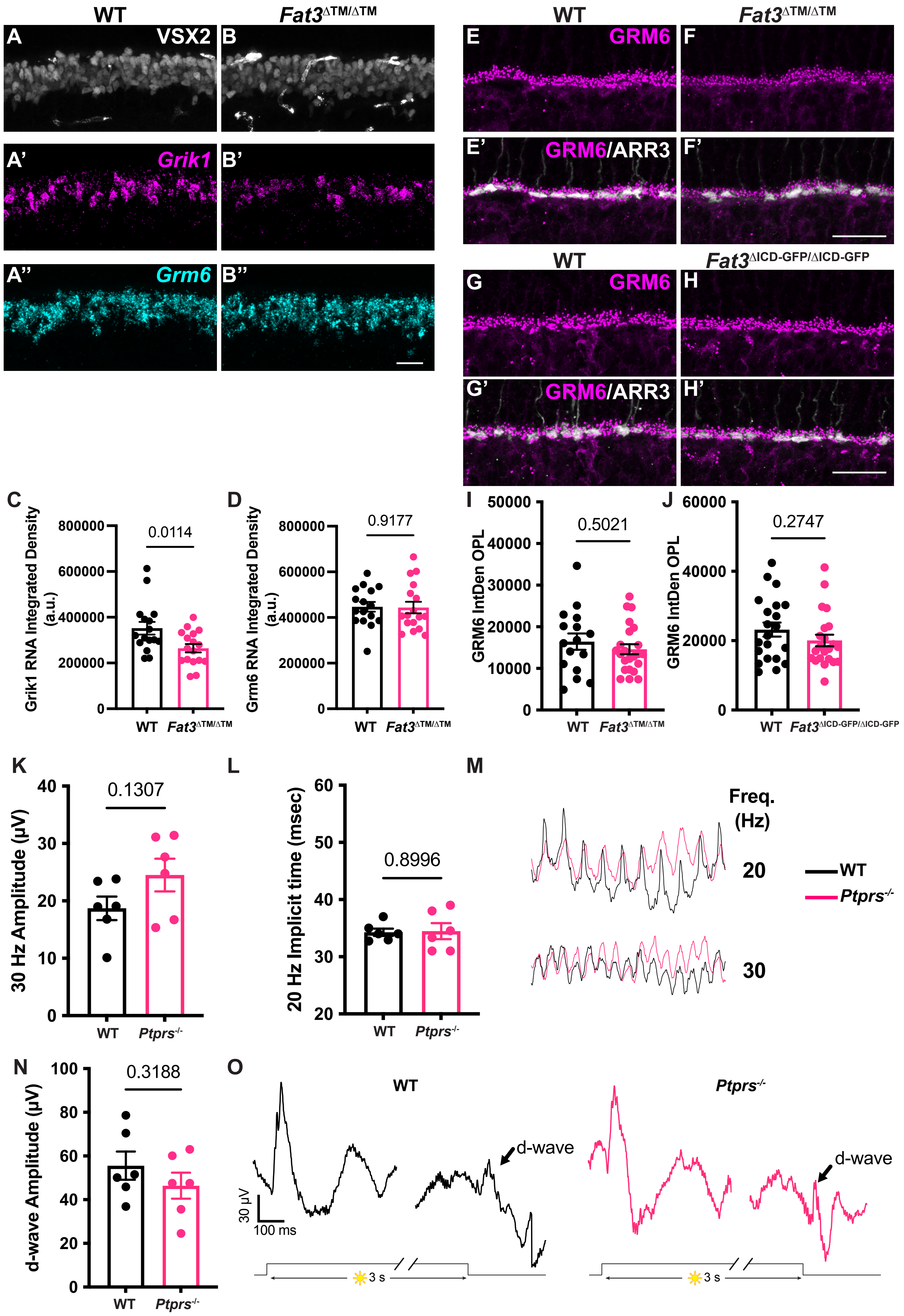
